# *Cis*-activation in the Notch signaling pathway

**DOI:** 10.1101/313171

**Authors:** Nagarajan Nandagopal, Leah A. Santat, Michael B. Elowitz

## Abstract

The Notch signaling pathway consists of transmembrane ligands and receptors that can interact both within the same cell (*cis*) and across cell boundaries (*trans*). Previous work has shown that *cis*-interactions act to inhibit productive signaling. Here, by analyzing Notch activation in single cells while controlling cell density and ligand expression level, we show that *cis*-ligands can in fact activate Notch receptors. This *cis*-activation process resembles *trans*-activation in its ligand level dependence, susceptibility to *cis*-inhibition, and sensitivity to Fringe modification. Cis-activation occurred for multiple ligand-receptor pairs, in diverse cell types, and affected survival and differentiation in neural stem cells. Finally, mathematical modeling shows how *cis*-activation could potentially expand the capabilities of Notch signaling, for example enabling “negative” signaling. These results establish *cis*-activation as a prevalent mode of signaling in the Notch pathway, and should contribute to a more complete understanding of how Notch signaling functions in developmental, physiological, and biomedical contexts.

## Introduction

The Notch signaling pathway enables intercellular communication in animals. It plays critical roles in diverse developmental and physiological processes, and is often mis-regulated in disease, including cancer (Louvi and Artavanis-Tsakonas 2012; Siebel and Lendahl 2017). Notch signaling occurs when membrane-bound ligands such as Dll1 and Dll4 on one cell activate Notch receptors on neighboring cells (Figure 1A, *trans*-activation) (Artavanis-Tsakonas, Rand, and Lake 1999; J. T. Nichols, Miyamoto, and Weinmaster 2007; Bray 2016). However, other types of interactions are also known to occur. Intercellular interactions between Notch1 and the ligand Jag1 have been shown to block *trans*-activation during angiogenesis and in cell culture (Figure 1A, *trans*-inhibition) (Benedito et al. 2009; Hicks et al. 2000; Golson et al. 2009). Additionally, Notch ligands and receptors co-expressed in the same cell have been shown to mutually inhibit one another, suppressing productive intercellular signaling (Figure 1A, *cis*-inhibition) (Sprinzak et al. 2010; del Álamo, Rouault, and Schweisguth 2011; Fiuza, Klein, and Martinez Arias 2010). Such ‘*cis*-inhibition’ has been shown to be important in diverse developmental processes including neurogenesis, wing margin formation in Drosophila, and maintenance of postnatal human epidermal stem cells (Micchelli, Rulifson, and Blair 1997; Jacobsen et al. 1998; Franklin et al. 1999; Lowell et al. 2000).

**Figure 1.**
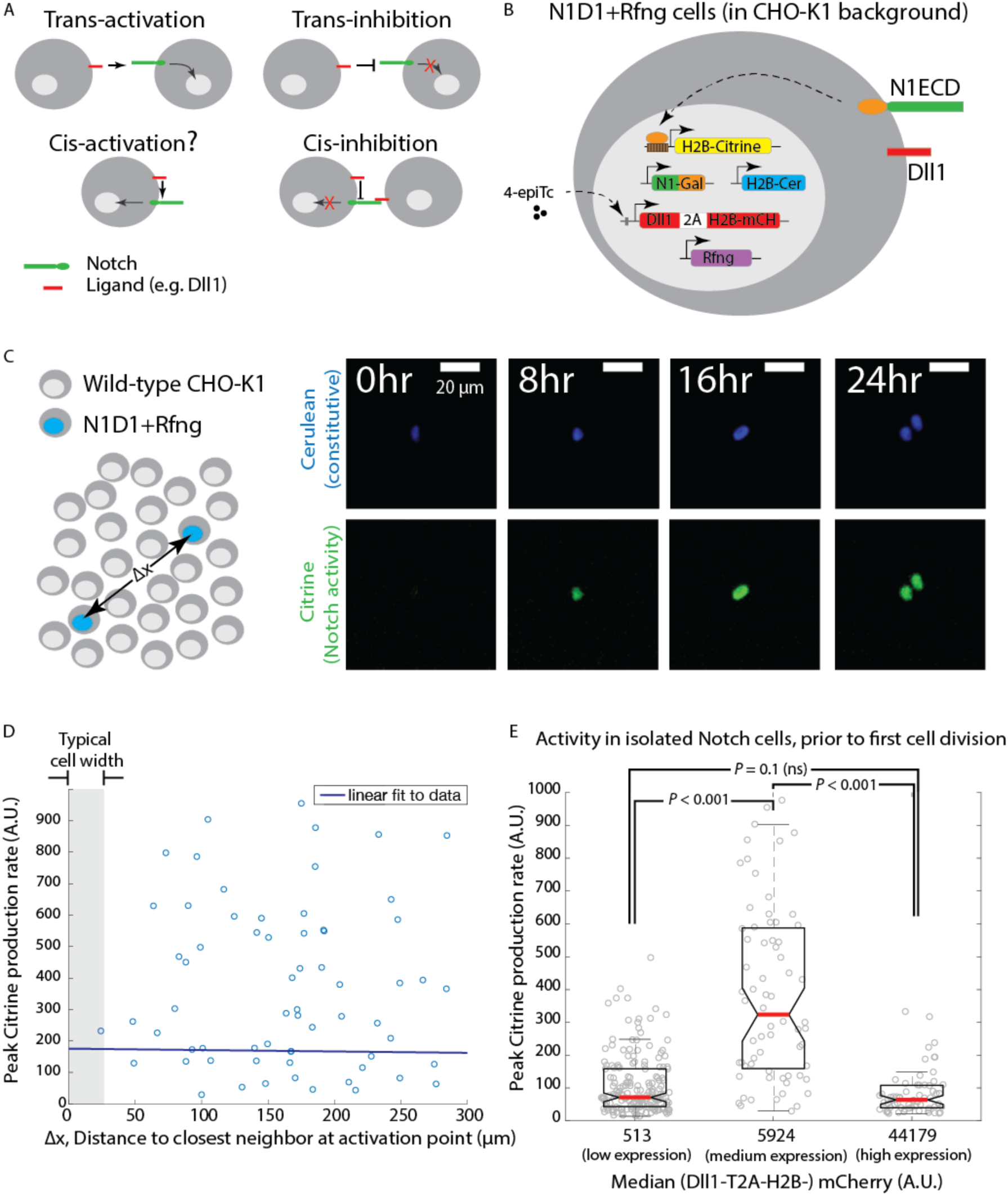
Engineered CHO-K1 N1D1+Rfng cells show ligand-dependent *cis*-activation. **(A)** Schematic of actual and potential *cis*- and *trans*-interaction modes in the Notch pathway. **(B)** Schematic of the N1D1+Rfng cell line. CHO-K1 cells were engineered to express a chimeric receptor combining the Notch1 extracellular domain (‘Notch1ECD’, green) with the Gal4 transcription factor (orange) in place of the endogenous intracellular domain. When activated, released Gal4 activates a stably integrated fluorescent H2B-Citrine reporter gene (yellow) through UAS sites (brown) on the promoter. Cells also contain a stably integrated construct expressing Dll1 (red) with a co-translational (2A, white) H2B-mCherry readout (‘mCH’, red), from a 4-epiTc-inducible promoter. Cells also constitutively express Rfng (purple) and H2B-Cerulean (‘H2B-Cer’, blue). **(C)** (*Left*) Schematic of *cis*-activation assay conditions. A minority of N1D1+Rfng (blue nuclei) cells were mixed with an excess of wild-type CHO-K1 cells (white nuclei). The typical distance between N1D1+Rfng cells is ∆x. (*Right*) Filmstrip showing activation (Citrine fluorescence, green) of an isolated N1D1+Rfng cell using time-lapse microscopy. Constitutive cerulean fluorescence (blue) in the same cell nucleus is also shown (see Video 1 for additional examples). **(D)** Peak Notch activation rate in isolated N1D1+Rfng cells (y-axis) versus distance to each of its closest neighboring N1D1+Rfng cell (x-axis) at the point of maximum activity. One cell width is indicated by gray shaded area. Solid blue line indicates linear fit, whose flat slope suggests a cell-autonomous, distance-independent process. **(E)** Box plots showing the distribution of peak Notch activation rates in isolated N1D1+Rfng cells prior to the first cell division in the *cis*-activation assay, for three different median Dll1 induction levels (see Figure 1 - figure supplement 2A for corresponding distributions). *P*-values calculated using two-sided KS-test.

The ability of co-expressed Notch ligands and receptors to interact on the same cell provokes the question of whether such interactions might also lead to pathway activation (Figure 1A, ‘*cis*-activation’). *Cis*-activation has been postulated (Formosa-Jordan and Ibañes 2014a; Hsieh and Lo 2012; Coumailleau et al. 2009; Pelullo et al. 2014), but has not been systematically investigated. A key challenge in identifying and characterizing such a behavior is the difficulty of discriminating between *trans*- and *cis*-activation in a multicellular tissue context, i.e. attributing any observed Notch signal to *trans* or *cis* ligand-receptor interactions. It has therefore remained unclear whether and where *cis*-activation occurs, how it compares to *trans*-activation, and how it might co-exist with *cis*-inhibition.

Here, we used single cell imaging to investigate activation in isolated cells. We find that *cis*-activation is a pervasive property of the Notch signaling pathway. It occurs for multiple ligands (Dll1 and Dll4) and receptors (Notch1 and Notch2), and in diverse cell types, including fibroblastic CHO-K1 cells, epithelial NMuMG and Caco-2 cells, and in neural stem cells. Cis-activation resembles *trans*-activation in the magnitude of the signaling response, modulation by R-Fringe, and susceptibility to *cis*-inhibition at high ligand concentrations. Furthermore, *cis*-activation appears to impact the survival and differentiation of neural stem cells. Finally, mathematical modeling shows that *cis*-activation could expand the capabilities of the Notch pathway, potentially enabling “negative” Notch signaling and integration of information about levels of *cis*- and *trans*-ligand. Together, these results extend the range of Notch signaling modes and provoke new questions about how *cis*-activation could function in diverse processes.

## Results

### Notch1-Dll1 cells show ligand-dependent cis-activation

To analyze *cis*-activation, we sought to develop a synthetic platform that could allow tuning of Notch pathway components and quantitative single-cell read-out of pathway activation (Figure 1B). We used the CHO-K1 cell line, which does not naturally express Notch receptors or ligands and has been used in previous studies of the Notch pathway (Sprinzak et al. 2010; LeBon et al. 2014; Nandagopal et al. 2018). We engineered these cells to co-express the Notch ligand Dll1, a chimeric Notch1ECD-Gal4 receptor, as well as the Gal4-activated H2B-Citrine fluorescent reporter gene that enables readout of Notch activation (Materials and methods). In these engineered cell lines, receptors are expressed constitutively. Dll1 expression can be induced using the small molecule 4-epi-Tetracycline (4-epiTc) in a dose-dependent manner, and monitored using a co-translational H2B-mCherry fluorescent protein (LeBon et al. 2014). Upon activation by Notch ligand, the chimeric N1ECD-Gal4 releases Gal4, which can travel to the nucleus and activate H2B-Citrine expression. Engineered cells also express a Radical Fringe (Rfng) gene, which enhances Notch1-Dll1 interactions through receptor glycosylation (Moloney et al. 2000). Finally, these ‘N1D1+Rfng’ cells also constitutively express nuclear-localized H2B-Cerulean fluorescent protein, which enables their identification in co-culture assays.

To discriminate *cis*-activation from *trans*-activation, we isolated individual N1D1+Rfng cells by co-culturing a minority of N1D1+Rfng cells (1%) with an excess of wild-type CHO-K1 cells (*‘cis*-activation assay’, Figure 1C, left). We first verified that their relative density was low enough to prevent *trans*-interactions between them, by confirming that a similar fraction of pure receiver cells, which express Notch1 but no ligands, were not activated by N1D1+Rfng cells (Figure 1-figure supplement 1). We then used time-lapse microscopy to measure Notch activity in N1D1+Rfng cells in the *cis*-activation assay (Materials and methods). At intermediate Dll1 expression levels (with 80 ng/ml 4-epiTc), isolated N1D1+Rfng cells showed clear activation (Figure 1C, right; Video 1). As expected for a cell-autonomous process, Notch activity, estimated by the peak rate of Citrine production, was uncorrelated with proximity to neighboring N1D1+Rfng cells (Figure 1D). However, the activity depended strongly on ligand expression levels (Figure 1E). Interestingly, this dependence was non-monotonic, peaking at intermediate levels of Dll1 induction, but returning to baseline at high ligand levels (Figure 1E, Dll1 induction levels shown in Figure 1-figure supplement 2A). This suppression of Notch activity is consistent with the previously described phenomenon of *cis*-inhibition (de Celis and Bray 1997; Sprinzak et al. 2010; del Álamo, Rouault, and Schweisguth 2011). These results suggest that Notch1 can be activated by intermediate concentrations of *cis*-Dll1, but that this *cis*-activation is dominated or replaced by *cis*-inhibition at high ligand concentrations.

We next asked how the strength of *cis*-activation compared to that of *trans*-activation, by analyzing the effect of intercellular contact on signaling levels. To control intercellular contact, we varied the fraction (relative density) of N1D1+Rfng cells in the co-culture, using wild-type CHO-K1 cells to maintain a constant total cell density. In order to increase the throughput of the experiment, we used flow cytometry to measure activation levels after 24 hours of culture (see Materials and methods). Total activation levels, which reflect a combination of *cis*- and *trans*-signaling, displayed a non-monotonic dependence on ligand expression for all N1D1+Rfng fractions, similar to *cis*-activation alone (Figure 1-figure supplement 2B, cf. Figure 1E). The peak amplitude of total activation was ~3-fold higher than *cis*-activation at high cell densities, but *cis*- and total signaling peaked at the same ligand concentration (Figure 1-figure supplement 2C). These results are consistent with overall Notch activation reflecting contributions from both *cis*- and *trans*-interactions, both of which depend similarly on ligand concentration

In principle, *cis*-activation could be an artifact of the chimeric Notch1ECD-Gal4 receptor. To test this possibility, we analyzed cells co-expressing Dll1 and the wild-type Notch1 receptor (N1^WT^). For readout, we used a previously characterized 12xCSL-H2B-Citrine reporter gene, which can be activated by cleaved NICD through multimerized CSL binding sites in the promoter region (Figure 1-figure supplement 3A, left panel) (Sprinzak et al. 2010). In the *cis*-activation assay, these ‘N1^WT^D1+Rfng’ cells showed *cis*-activation and non-monotonic dependence on ligand levels, similar to the responses described above for the N1ECD-Gal4 cells (Figure 1-figure supplement 3A, right panel). These results indicate that *cis*-activation occurs for wild-type as well as engineered receptors.

Next, we asked whether *cis*-activation occurs in other cell types. We turned to the mammary epithelial cell line NMuMG (Owens, Smith, and Hackett 1974), knocking out the endogenously expressed Notch2 and Jagged1 genes using CRISPR-Cas9 (Materials and methods, Figure 1-figure supplement 3B), and adding an inducible Dll1, constitutive Rfng, and a Notch1 reporter system similar to the one used in our CHO-K1 cells, but with a different Dll1 induction system (Figure 1-figure supplement 3C). To ensure proper apical localization in these polarized cells, we also attached the ankyrin (ANK) domain of NICD to the Notch1ECD-Gal4 protein (Materials and methods, Figure 1-figure supplement 3D). When analyzed in the *cis*-activation assay, isolated NMuMG N1D1+Rfng cells showed Notch activation (Figure 1-figure supplement 3E, Video 2). This activation increased with ligand expression in a dose-dependent manner (Figure 1- figure supplement 3F), indicating that epithelial NMuMG cells also display *cis*-activation.

To test whether *cis*-activation occurs with endogenous ligands and receptors, we next analyzed a human colorectal adenocarcinoma cell line Caco-2, in which Notch signaling is known to regulate proliferation and differentiation (Sääf et al. 2007; Dahan et al. 2011). We measured endogenous Notch activity in these cells by transfecting them with the 12xCSL-H2B-Citrine reporter construct (used in Figure 1-figure supplement 3A). To analyze *cis*-activation, we plated the transfected cells sparsely, with or without treatment with the Notch inhibitor DAPT (Dovey et al. 2001) (see Materials and Methods). After 24 hours, cells treated with DAPT displayed lower levels of Notch activation compared to untreated cells, suggesting that these cells can *cis*-activate (Figure 1-figure supplement 4A). The level of *cis*-activation appeared similar to activation levels in cells plated at high density (*cis* + *trans*) (Figure 1-figure supplement 4B, Materials and Methods), indicating that the magnitude of *cis*-signaling is comparable to that shown at higher densities.

Taken together, our results demonstrate that *cis*-activation is a general phenomenon in Notch signaling, occurring in diverse cell types. In cells co-expressing Notch1, Dll1, and R-Fringe, *cis*-activation strength depends on ligand concentration, and is non-monotonic in CHO cells, where *cis*-activation peaks at intermediate ligand concentrations, but is replaced by *cis*-inhibition at the highest ligand levels (Figure 1-figure supplement 2A, C).

### Cis-activation changes with ligand-receptor affinity

The affinity of ligand-receptor interactions is affected by the identities of the ligand and receptor, and co-expression of glycosyltransferases like Rfng (Moloney et al. 2000; Yang et al. 2005; Taylor et al. 2014). We next asked how these factors impact *cis*-activation, starting with Rfng, which is known to increase Notch1-Dll1 signaling. We compared the N1D1+Rfng line to its parental line (‘N1D1’), which lacks ectopic Rfng but is otherwise identical. N1D1 cells also showed ligand-dependent *cis*-activation, but at reduced levels (Figure 2A). Non-monotonicity was preserved, with *cis*-activation dominating at intermediate Dll1 concentrations, and *cis*-inhibition dominating at high Dll1 concentrations. Further, extending the analysis of Notch activation to increased intercellular contact, we observed a similar dependence on cell fraction and Dll1 expression with and without Rfng, with the two states differing in signal amplitude but not the shape of the ligand response (Figure 2B). These results show that Rfng increases the amplitude of both *cis* and *trans* signaling without affecting the overall dependence of signaling on Dll1 expression level.

**Figure 2.**
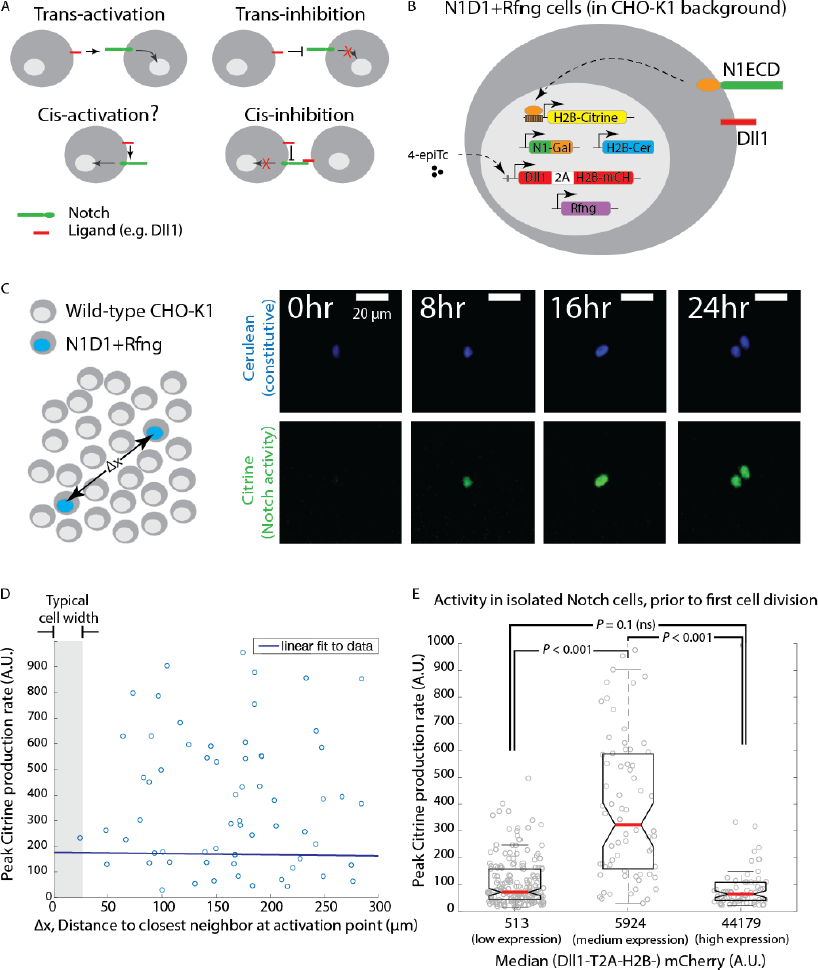
Cis*-activation is affected by changes in ligand-receptor affinity*. **(A)** (*Top*) Cell lines used for analyzing effect of Rfng on *cis*-activation. (*Bottom*) Plots showing mean Notch activation (reporter Citrine fluorescence normalized to background fluorescence in uninduced cells) in N1D1 (black) or N1D1+Rfng (purple) cells expressing different levels of Dll1 (measured using co-translational mCherry fluorescence). Error bars indicate s.e.m (n = 3 biological replicate experiments). **(B)** Heatmaps of mean Notch activation (n = 3 biological replicates), relative to background reporter fluorescence, in N1D1+Rfng (upper panel) or N1D1 (lower panel) cells induced with different [4-epiTc] (columns) and cultured at different relative fractions (rows). Upper panel is the same data in Figure 1-figure supplement 2B, re-plotted for direct comparison. In both cell lines, Rfng expression predominantly affects signal amplitude (compare intensity scales between heatmaps). **(C,D)** (*Top*) Cell lines used for analyzing effect of ligand on *cis*-activation of Notch1 (C) or Notch2 (D). (*Bottom*) Comparison of mean *cis*-activation in polyclonal populations (‘Pop’) of cells co-expressing Dll1 or the higher affinity ligand Dll4 with the indicated receptor, as a function of ligand expression, read out by co-translated H2B-mCherry fluorescence. Values represent mean of 3 biological replicates. Error bars indicate s.e.m. Note difference in y-axis scales between panels C and D.

We next analyzed how the identity of the ligand affects *cis*-activation. Compared to Dll1, the ligand Dll4 has increased affinity for Notch1 (Andrawes et al. 2013). We engineered CHO-K1 cells to stably express either an inducible Dll4-T2A-H2B-mCherry or Dll1-T2A-H2B-mCherry, along with a constitutive Notch1ECD-Gal4 Notch reporter system. To enable direct comparison, we performed the *cis*-activation analysis on polyclonal populations for the two cell lines. Compared to the Dll1-expressing cells, Dll4-expressing cells showed enhanced *cis*-activation and *cis*-inhibition, exhibiting greater peak reporter activity at intermediate ligand expression levels but comparable activity at the highest ligand expression levels (Figure 2C). Adding Rfng to the N1D4 cells did not further increase *cis*-activation or *cis*-inhibition (Figure 2-figure supplement 1A), consistent with the idea that Rfng does not increase Dll4-Notch1 affinity (Taylor et al. 2014). Together, these data suggest that stronger ligand-receptor interactions, achieved either through the addition of Rfng or through the use of a stronger affinity Notch ligand like Dll4, can enhance both *cis*-activation and inhibition.

### Notch2 shows stronger *cis*-activation but decreased *cis*-inhibition compared to Notch1

To investigate whether *cis*-activation occurs with other Notch receptors, we engineered CHO-K1 cells to express a similar reporter system for Notch2 activation (Notch2ECD-Gal4) along with inducible Dll1- or Dll4-T2A-H2B-mCherry, as described previously. Both N2D1 and N2D4 cell populations showed a notable increase in *cis*-activation compared to their Notch1 counterparts, with ~3-fold higher maximal *cis*-activation (Figure 2D, note difference in scale compared to Figure 2C). Moreover, unlike Notch1, Notch2 showed similar levels of *cis*-activation by the Dll1 and Dll4 ligands (Figure 2D). Strikingly, the profile of activation was monotonic, with *cis*-activation persisting even at the highest ligand levels tested (Figure 2-figure supplement 2). Together, these results indicate that Notch2 undergoes *cis*-activation, does so at a higher level than Notch1, and is not cis-inhibited as strongly as Notch1.

### Cis-activation regulates neural stem cell maintenance and differentiation

We next asked whether *cis*-activation could play a functional role in Notch-mediated processes. As a model system, we used mouse cortical neural stem cells (NSCs), in which Notch signaling is known to regulate self-renewal and differentiation (Bertrand, Castro, and Guillemot 2002; Kageyama et al. 2008). Importantly, primary NSCs can be cultured and propagated *in vitro* under defined media conditions and cell density (Daadi 2002).

To assess their potential for cis-activation, we first sought to identify the Notch components expressed in NSCs. Bulk RNA sequencing showed that these cells express high levels of the receptor Notch1, the ligand Dll1, and the Lfng glycosyltransferase, and lower levels of Notch2 and Rfng, suggesting that NSCs have the potential to *cis*-activate (Figure 3-figure supplement 1A, Materials and methods). To identify suitable gene targets for assaying Notch activation, we next analyzed the expression of the *Hes/Hey* genes, with or without treatment with the Notch inhibitor DAPT for 12 hours.

Since NSC culture conditions include treatment with the EGF and FGF growth factors, and there is evidence for crosstalk between the growth factors and Notch signaling pathways in these cells (Aguirre, Rubio, and Gallo 2010; Nagao, Sugimori, and Nakafuku 2007), we compared Notch activation with or without the Notch inhibitor DAPT (10μM), under both reduced (0.5 ng/ml EGF, no FGF) and normal (20 ng/ml EGF, 20 ng/ml FGF) growth factor conditions (Materials and methods). Canonical Notch target genes *Hes1*, *Hes5*, and *Hey1* responded to Notch inhibition in these cells, and did so most strongly at reduced growth factor concentrations (Figure 3-figure supplement 1B).

To analyze *cis*-activation in NSCs, we plated them at very low density in reduced growth factor conditions (0.1 ng/ml EGF, no FGF), and cultured them with or without 10 μM DAPT (Figure 3A, Materials and methods). After 6 hours, we assayed mRNA transcript levels of Hes1, Hey1, and Hes5 in isolated cells using single-molecule HCR-FISH (Choi et al. 2010, 2018) (Figure 3B, Figure 3-figure supplement 1C). Consistent with Notch-induced gene expression in these isolated cells, DAPT treatment decreased mean expression levels of all three genes (Figure 3B, cumulative histograms). Specifically, the expression levels decreased by ~2-fold for Hes1 (mean fold change 2.5, bootstrapped 95% confidence interval [2.1, 4.1]), and Hes5 (mean 1.9, c.i. [1.3, 2.5]), and by ~25% for Hey1 (1.2, c.i. [1.1, 1.2]). These results suggest that isolated NSCs show Notch-dependent gene expression, consistent with *cis*-activation.

**Figure 3.**
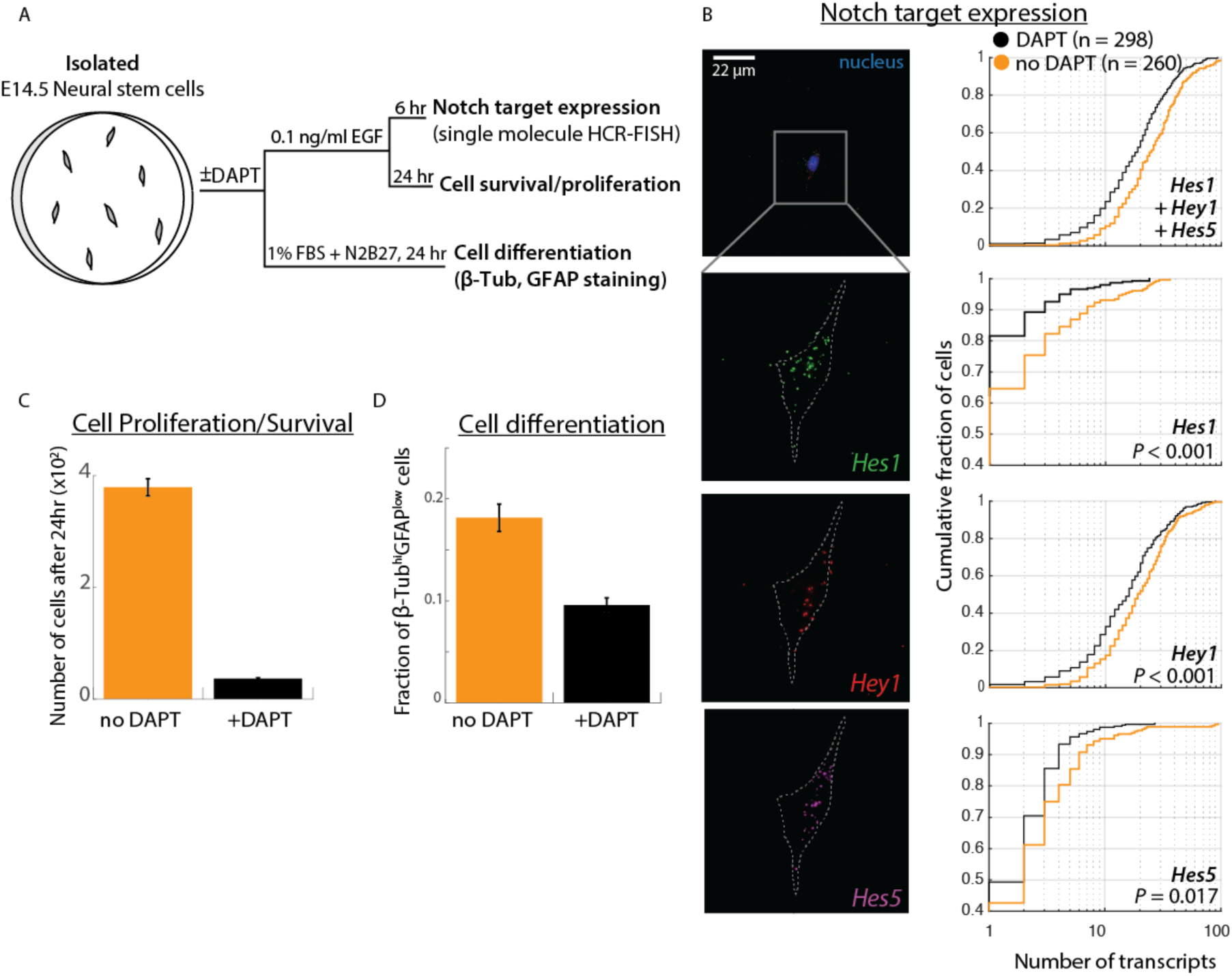
Cis-activation occurs in neural stem cells and regulates survival and differentiation. **(A)** E14.5 mouse cortical neural stem cells (NSCs) were plated sparsely and treated with ±10 μM DAPT, cultured under growth or differentiation conditions, and subsequently assayed for expression of Notch target genes, survival, or expression of differentiation markers. **(B)** (*Left*) Representative example of an isolated NSC (top panel, DAPI-stained nucleus shown; note lack of neighboring cells within ~50 μm) not been treated with DAPT, assayed for expression of Hes1 (green), Hey1 (red), and Hes5 (magenta) mRNA using multiplexed single-molecule HCR-FISH (see Materials and methods). (*Right*) Cumulative distribution plots of gene expression in DAPT-treated (black) and untreated (orange) cells (see Materials and methods for transcript quantification). *P*-values calculated using two-sided KS-test. See Figure 3-figure supplement 1C for additional examples of isolated cells showing Hes/Hey expression. **(C)** Comparison of mean number of isolated untreated (orange) or DAPT-treated (black) cells after 24 hours. Error bars represent s.e.m of n=4 biological replicates. **(D)** Fraction of cells showing high β-Tubulin + low GFAP levels, based on immunostaining (see Materials and methods) under differentiation conditions. Error bars represent s.e.m, n=2 biological replicates. See Figure 3-figure supplement 2B for complete distributions.

We next asked whether *cis*-activation plays a functional role in regulating the Notch-dependent processes of NSC maintenance and differentiation. To test the effect on cell maintenance, we plated cells at low density in low growth factor conditions with or without DAPT treatment (see Materials and methods). Under these conditions, DAPT treatment dramatically decreased the number of cells after 24 hours, implying that initial cell survival depends on *cis*-activation (10-fold, Figure 3C). Moreover, plating NSCs on recombinant Dll1ext-IgG led to a striking increase in cell numbers, further supporting a positive effect of Notch activation on cell survival (Figure 3-figure supplement 2A, Materials and methods). Together, these results suggest that *cis*-activation of Notch is necessary for initial NSC survival in low growth factor conditions.

To assay the effect of *cis*-activation on NSC differentiation, we cultured NSCs at low density in low serum media previously shown to induce differentiation into neural and glial fates (Imayoshi et al. 2013), with or without DAPT treatment. We estimated differentiation by immunostaining for the marker genes β-Tubulin and GFAP after 24 hours. Quantification of staining in single cells showed that DAPT treatment altered expression patterns of these markers, suggesting that *cis*-activation could influence cell fate choice as well as survival (Figure 3D, Figure 3-figure supplement 2B, Materials and methods).

### Cis-activation requires cell surface interactions between ligands and receptors

To gain insight into where *cis*-activation occurs in the cell, we tested whether cell surface ligand-receptor interactions were required for productive signaling (Figure 4A). Treatment of cells with soluble recombinant N1ECD-Fc (rN1ECD-Fc) receptors has been shown to prevent *trans*-signaling by blocking surface ligands (Klose et al. 2015). We first confirmed that activation levels in densely-plated N1D1+Rfng cells decreased when they were incubated in rN1ECD-Fc containing media for 24 hours, compared to IgG-treated controls (Figure 4-figure supplement 1A). Interestingly, a similar decrease in activation levels could be observed in N1D1+Rfng cells plated in the *cis*-activation assay (Figure 4B), suggesting that blocking ligand-receptor interactions at the surface reduces *cis*-activation. This effect was not limited to soluble receptor fragments; co-culturing a minority (5%) of N1D1+Rfng cells with an excess of Notch1-only expressing cells similarly reduced *cis*-activation, and by a comparable amount (Figure 4-figure supplement 1B). We further perturbed cell-surface ligand-receptor interactions by treating cells with Blebbistatin, an inhibitor of non-muscle myosin II, known to disrupt cellular adhesion and protrusions (Materials and methods) (Liu et al. 2010; Shutova et al. 2012). Similar to rN1ECD-Fc, treatment with Blebbistatin decreased both *cis*- and *trans*-activation of N1D1+Rfng cells to similar extents (Figure 4-figure supplement 1C). These results suggest that *cis*-activation requires surface presentation of the ligand, and is consistent with a model in which productive *cis* ligand-receptor interactions, like *trans* interactions, occur at the cell surface.

**Figure 4.**
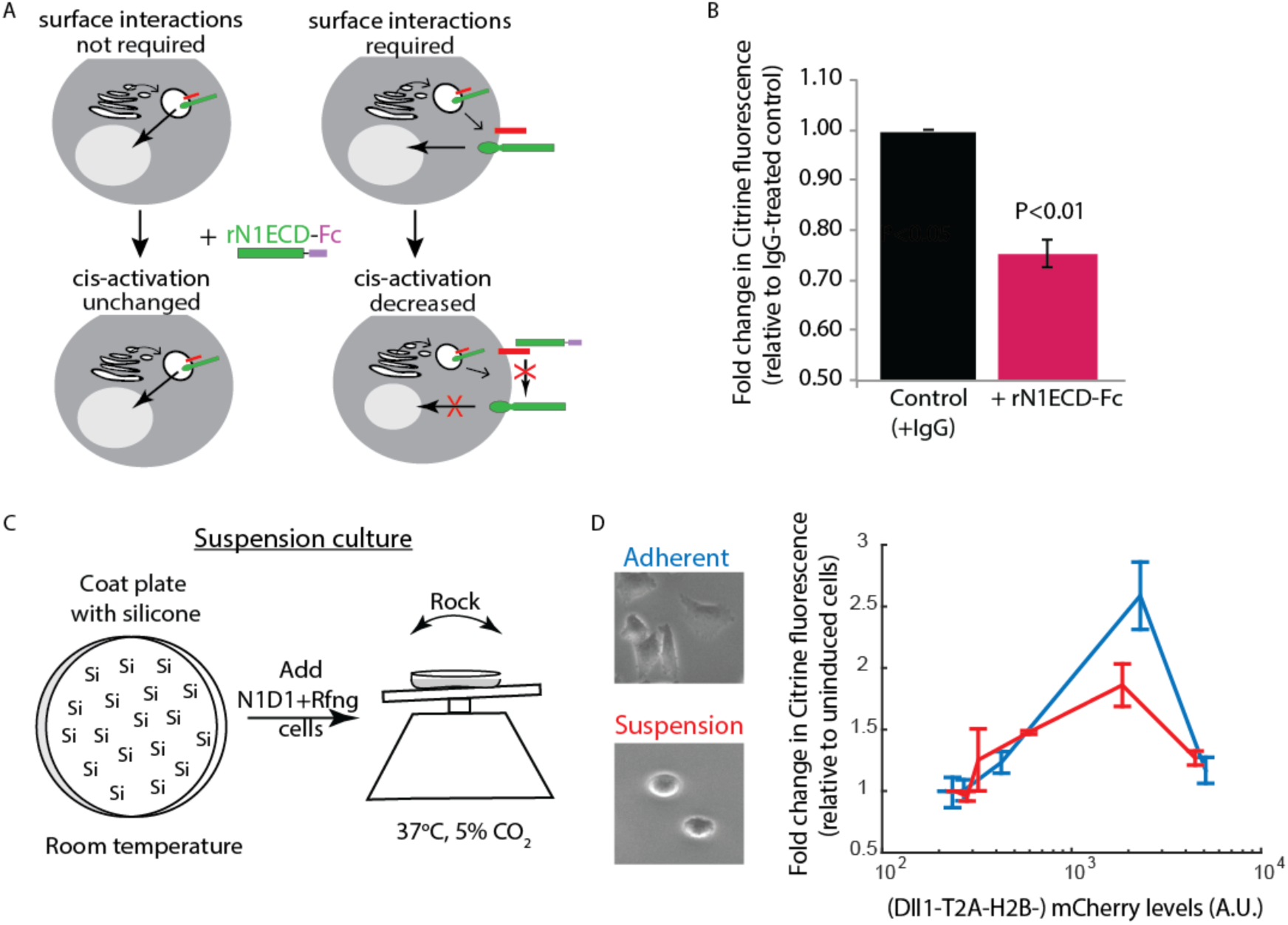
Receptor-ligand cell surface interactions are necessary for *cis*-activation. **(A)** Schematics showing how soluble recombinant N1ECD-Fc protein (rN1ECD-Fc) can be used to test whether surface interactions between ligand (red) and receptor (green) are necessary for *cis*-activation. (*Left*) With intracellular *cis*-activation, addition of extracellular rN1ECD-Fc should not affect *cis*-activation levels. (*Right*) If surface interactions are necessary for *cis*-activation, rN1ECD-Fc treatment should reduce activation levels by competing with receptors for cell-surface ligands. **(B)** Comparison of mean Notch activation in N1D1+Rfng cells incubated with rNotch1ECD-Fc receptors (magenta) or an IgG control (black) for 24 hours (see Materials and methods). Cells were plated for a *cis*-activation assay and analyzed by flow cytometry <24 hours post-plating. Error bars represent s.e.m (n=3 biological replicate experiments). *P*-values calculated using the one-sided Student T-test. **(C)** Schematic of procedure for culturing N1D1+Rfng cells in suspension. Plates were coated with a Silicone solution (‘Si’), and cells were subsequently plated for a *cis*-activation assay (co-culture of 5×10^3^ N1D1+Rfng + 150×10^3^ CHO-K1 cells). The plate was incubated at 37°C, 5% CO_2_ on a rocker to prevent cells from adhering to the plate surface. **(D)** (*Left*) Representative image of cells grown in adherent (blue) or suspension (red) conditions. (*Right*) Comparison of mean Notch activation levels, relative to background reporter fluorescence, in N1D1+Rfng cells cultured for 24 hours in suspension (red) or in adherent conditions (blue), for different Dll1 expression levels (measured using co-translated mCherry fluorescence). Cells were in *cis*-activation co-culture conditions (5×10^3^ N1D1+Rfng + 150×10^3^ CHO-K1 cells). Error bars represent s.e.m (n = 3 biological replicates).

To determine whether interactions other than surface receptor/ligand binding could be facilitating *cis*-activation, we sought to measure the ability of N1D1+Rfng cells to *cis*- activate in a suspension culture. This enabled us to ask whether cis-activation requires interactions with the culture dish surface. For example, *cis*-activation could require focal adhesions formed at points of contact with the dish or result from cells depositing ligands on the culture surface, which could in turn *trans*-activate cell-surface receptors. To create suspension cultures, cell adhesion to the plate surface was prevented by pre-coating the surface with silicone (Nienow, Hewitt, and Heathman 2016) and putting the plate on a rocker for the duration of the experiment (Figure 4C, Materials and methods). Under such suspension culture conditions, N1D1+Rfng cells (co-cultured with an excess of wild-type CHO-K1 cells as in the *cis*-activation assay, see Figure 1C) continued to show *cis*-activation (Figure 4D). We note that culturing cells in suspension leads to slight reductions in cell-surface Notch1 and Dll1 levels (Figure 4-supplement 1D), which could account for the minor reduction in peak *cis*-activation observed in suspension cells compared to adherent cells. Nevertheless, these results suggest that *cis*-activation is a cell-intrinsic process and does not require extensive interactions with the culture surface.

Together, our results support a model in which *cis*-activation arises in a cell-intrinsic manner from interactions between ligands and receptors on the cell surface. More generally, the observed similarities between *cis*- and *trans*-activation in their dependence on ligand concentration and ligand-receptor affinities, and sensitivity to perturbations, could reflect a common underlying mechanism of activation.

### *Cis*-activation enables integration of intra- and extracellular information and negative signaling

Finally, we turned to the fundamental question of what types of underlying interactions could explain the observed characteristics of *cis*-activation, such as its non-monotonic dependence on ligand levels. We developed a series of simplified mathematical models of Notch-ligand *cis*-interactions at steady-state and compared their behaviors to experimental observations.

We first considered the simplest case of Notch (N) and Delta (D) reversibly interacting in *cis* to form a single activation-competent complex, denoted *C^+^*, which can subsequently undergo cleavage to release NICD (Figure 5A, Model 0, Materials and methods). We simulated this model using 10,000 biochemical parameter sets, chosen using the Latin Hypercube Sampling method (McKay, Beckman, and Conover 1979) (Figure 5B, Figure 5-figure supplement 1, Materials and methods). For each parameter set, we quantified the degree of non-monotonicity in the concentration of C^+^ as a function of the Delta production rate, α_D_ (Figure 5B, D). Model 0 did not produce non-monotonic responses (Figure 5C, Materials and methods), indicating that *cis*-activation at low ligand concentrations and *cis*-inhibition at high concentrations cannot both result from a single underlying type of *cis*-complex.

**Figure 5.**
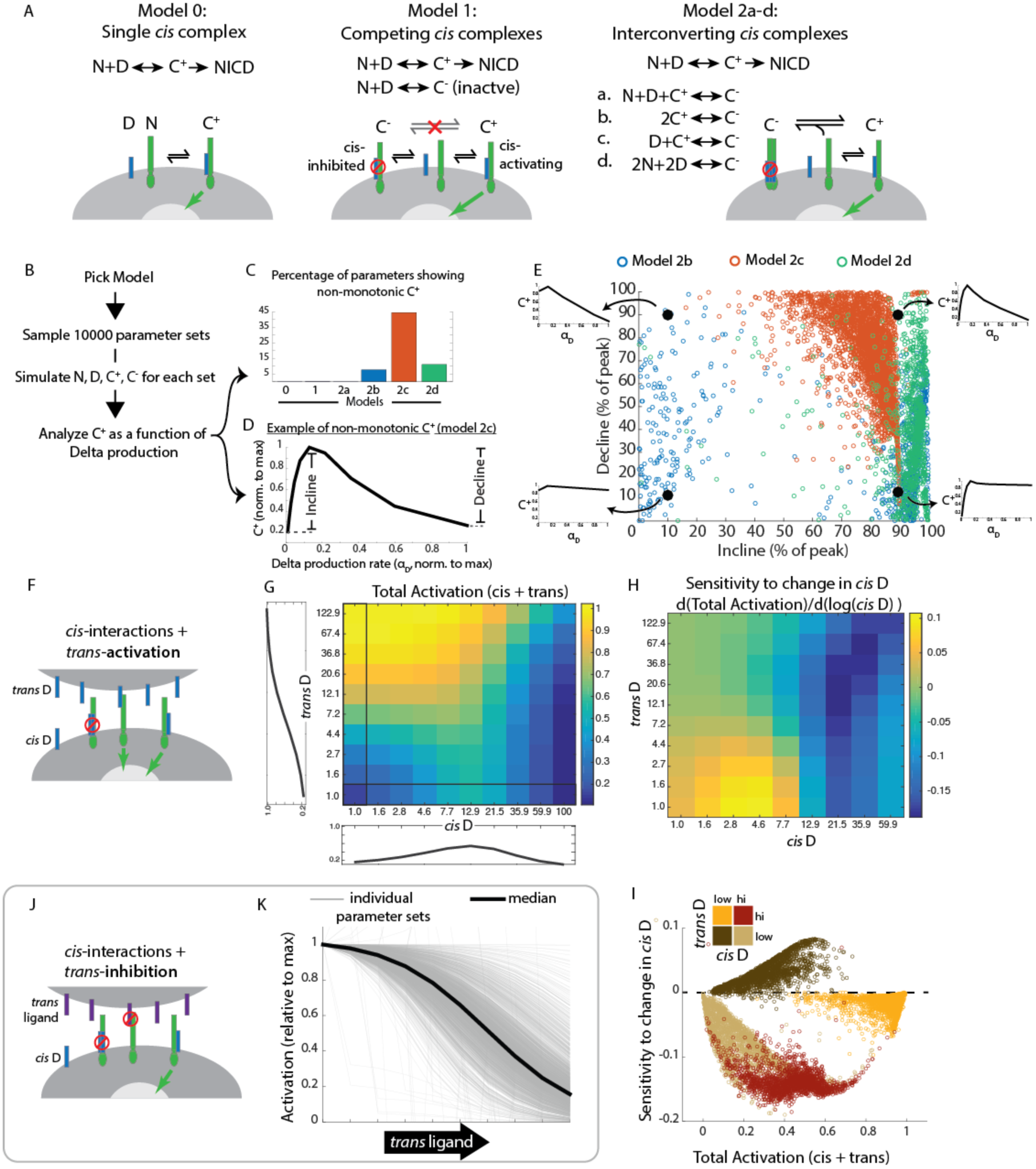
Mathematical model of *cis*-activation reveals potential roles in signal processing. **(A)** Schematics of different cis-activation models. In each model, Notch (‘N’, green) and Delta (‘D’ blue) interact to produce one or more cis-complexes, which can be active (‘C^+^’), producing NICD (green arrow) or inhibited (‘C^−^’, red circle). In Models 2a-d, C^+^ is formed through the same interaction but C^−^ formation is different for each of the included models. **(B)** Overview of simulations (se Materials and methods). **(C-E)** Results of simulations. **(C)** Percentage of parameters that lead to non monotonic C^+^ behavior in each of the models (see Material and methods for assessment of this feature) **(D)** Example of non-monotonic dependence of C^+^ on Delta production rate (‘α_D_’), generated in Model 2c for one choice of parameter values. The fractional incline and decline features used to characterize the degree of non-monotonicity (and plotted in panel E) are shown. **(E)** Scatter plot of fractional incline vs. decline for each non-monotonic C^+^ profile produced by Models 2b-d. Filled black circles and associated schematic plots highlight C^+^ profile shapes corresponding to different incline vs decline levels. **(F)** Schematic of model including both *cis*- and *trans*-activation. Notch receptors ca interact with intracellular Delta (‘*cis* D’) or extracellular Delta (‘*trans* D’) to form *cis*- and *trans* complexes, respectively. Cis-complexes can be either inhibited (red circle) or activating (green arrow) while *trans* complexes are activating. **(G)** Example of total activation (levels of activating *cis* + *trans* complexes) as a function of *cis* and *trans* D, for a single set of parameters producing non-monotonic *cis*-activation. **(H)** Corresponding sensitivity to change in *cis* D for the example in G. This sensitivity (‘d(Total Activation)/d(log(cis D))’) is defined as the change in total activation upon constant fold changes in *cis* D levels, and is derived from G by computing the difference between adjacent columns of the total activation matrix. **(I)** Scatter plot showing median values of total activation vs. sensitivity t change in *cis* D in different regimes of *cis* and *trans* D (high *cis*/high *trans* - red, high *cis*/low *trans* beige, low *cis*/high *trans* - orange, low *cis*/low *trans* - brown). Each circle represents results obtained using a single set of parameters in Model 2c (with *trans*-activation). **(J)** Schematic of model including *cis*- and *trans*-inhibition. Notch receptors can interact with intracellular Delta (‘*cis* D’, blue) o extracellular ligand (‘*trans* ligand’, purple) to form *cis*- and *trans*-complexes, respectively. Cis complexes can be either inhibited (red circle) or activating (green arrow), while *trans* complexes cannot activate. **(K)** Dependence of total activation levels on *trans*-ligand, for *cis* D production rate corresponding to peak *cis*-activation. Each grey line represents behavior for a single set of parameters, while the black line represents the median response across all tested parameters Simulation code and parameter values are included in Figure 5 – source data 1.

We next considered a more complex model in which Notch and Delta can generate two distinct types of cis-complexes, C^+^ and C^−^, with only the former competent to activate (Figure 5A, Model 1, Materials and methods). However, analysis across diverse parameter sets showed that this model too was unable to recapitulate non-monotonic C^+^ behavior (Figure 5C, Materials and methods).

We reasoned that preferential formation of active C^+^ and inactive C^−^ at lower and higher Delta production rates, respectively, could in principle result from a sequential process of *cis*-complex formation (Figure 5A). For instance, C^−^ could be formed from *C^+^* through additional interactions of the active complex with itself (C^+^ + C^+^ → C^−^, Model 2b), through interactions with free N and D (N + D + C^+^ → C^−^ Model 2a), or interactions with ligand (D+ C^+^ → C^−^, Model 2c). (Note that C^−^ has different constituents in each of these different models). Alternatively, formation of C^−^ might require higher-order N-D interactions than required for formation of C^+^. For example, C^−^ could require interaction of 2 ligands and 2 receptors (2N + 2D → C^−^, Model 2d). Notch ligands and receptors are known to form clusters (Bardot et al. 2005; James T. Nichols et al. 2007; Nandagopal et al. 2018), and each of these models is consistent with a picture in which the inactive C^−^ complex involves increased clustering of ligands and receptors compared to the active C^+^ complex. Strikingly, across the same set of parameters tested before, these models produced non-monotonic C^+^ profiles more frequently than Models 0 and 1 (Figure 5C). For example, Model 2c gave rise to non-monotonic behavior for over 40% of tested parameter sets. Moreover, when the shape of the C^+^ profile was analyzed across parameter sets that produced non-monotonicity (Figure 5D), Model 2c came closest to reproducing the experimentally observed cis-activation response shape, often showing nearly complete attenuation of activation at the highest Delta production rates (Figure 5E, cf. Figure 1E, Figure 1-figure supplement 2C). Other models often produced non-monotonic C^+^ profiles with more modest declines of 30-40% at the highest ligand production rates. While this analysis does not uniquely identify a specific molecular mechanism, it suggests that multiple distinct *cis*-complexes are likely required to explain the observed non-monotonic responses and that this behavior could arise through several distinct modes of interaction. It is also consistent with a potential role for a clustering-based mechanism.

In natural tissue contexts, *cis*-interactions co-exist with *trans*-interactions. To understand how these two effects combine, we extended Model 2c to incorporate the effects of *trans* interactions. We assumed that *trans*-ligands interact with Notch to form productive *trans* complexes, denoted T, and do so with the same rates of formation, dissociation, and degradation as the active *cis*-complexes, C^+^ (Figure 5F). For each non-monotonic parameter set in Figure 5E, we quantified the total concentration of active complexes (T+ C^+^) across a range of *trans*-Delta levels and *cis*-Delta production rates (see Materials and methods). Interestingly, as shown in Figure 5G for one parameter set, levels of activation were typically similar in different regimes of the *cis*-Delta and *trans*-Delta matrix. For example, medium *cis*-Delta + low *trans*-Delta and low *cis*-Delta + medium *trans*-Delta regimes (left middle and bottom middle, Figure 5G) showed higher but similar activation. In this model, the cell would thus be unable to distinguish between different combinations of *cis* and *trans* Delta based only on the total levels of activation. By contrast, the sensitivity of Notch activation to changes in *cis*-Delta exhibited a striking difference between these otherwise similar signaling regimes (Figure 5H). A more comprehensive analysis of sensitivity to 1.6-fold changes in *cis*-Delta showed that regimes of *cis*-Delta*/trans*-Delta with similar levels of Notch activation exhibited distinct sensitivities to changes in ligand expression level (Figure 5I). This analysis suggests that, in principle, a cell could individually estimate the levels of *cis* and *trans* ligand by comparing Notch activation levels before and after a change in *cis*-Delta expression. It will be interesting to see if such a dynamic mechanism is utilized naturally.

Finally, we asked whether *cis*-activation, by establishing an elevated basal level of Notch activity, could expand the kinds of *trans*-signaling modes that are possible in the Notch system. Jagged1 has been shown to form inactive *trans* complexes with Notch receptors glycosylated by Lfng (Shimizu et al. 2001; Moloney et al. 2000; LeBon et al. 2014). To test the effects of such *trans*-inhibitory interactions in the context of *cis*-activation, we extended Model 2c by incorporating an inactive *trans* complex, T-, and analyzed the dependence of Notch activity on the concentration of *trans* ligand (Figure 5J, K, Materials and methods). Interestingly, at *cis*-Delta expression levels that produce peak *cis*-activation, the presence of an inhibitory *trans*-ligand led to strong, dose-dependent decreases in Notch activity across multiple parameter sets (Figure 5K). This effect results from *trans*-ligands effectively removing the pool of Notch receptors available to *cis*-activate. In this way, *cis*-activation could enable a ‘negative’ mode of intercellular Notch signaling, where exposure to unproductive or weak extracellular ligands reduces signal in the cell. This negative mode of Notch signaling would complement the standard activating mode in much the same way that repressors complement activators in gene regulation.

## Discussion

*Cis*-activation is an example of autocrine signaling, which occurs in cytokine, Wnt, BMP, and other signaling pathways (Fang et al. 2013; Feinerman et al. 2010; Babb et al. 2017; Shukunami et al. 2000; Yokoyama et al. 2017). Typically, autocrine signaling occurs when molecules (e.g. hormones) released from a cell bind to and activate receptors on the cell from which they were synthesized (Leibiger et al. 2012). However, autocrine signaling can also occur from membrane anchored molecules located on the same cell. For example, *cis*-activation of cell adhesion molecules (CAMs) help induce neurite outgrowth during neuronal development (Sonderegger and Rathjen 1992).

While autocrine signaling occurs in other pathways, it has only been minimally explored in the Notch pathway (Formosa-Jordan and Ibañes 2014b; Hsieh and Lo 2012). One reason for this may be that the previously characterized inhibitory cis-interactions could appear to rule out cis-activation (Sprinzak et al. 2010; LeBon et al. 2014; Miller, Lyons, and Herman 2009). Additionally, it can be difficult to disentangle *cis*- and *trans*-activation in the context of a tissue or an *in vitro* system where *cis*-ligands and *trans*-ligands are simultaneously present (Eddison, Le Roux, and Lewis 2000; Hartman, Reh, and Bermingham-McDonogh 2010; Daudet and Lewis 2005; Sprinzak et al. 2010). To address these issues, we analyzed *cis*-activation in spatially isolated cells engineered with a Notch reporter system that allows tunable control of ligand expression. This assay demonstrated *cis*-activation of the receptors Notch1 and Notch2 by the ligands Dll1 and Dll4. We also detected *cis*-activation across multiple cell types using both engineered and endogenous Notch components, suggesting that *cis*-activation is a general phenomenon in Notch signaling. Strikingly, our study not only revealed that *cis*-activation can co-exist with *trans*-activation, but showed that *cis*-activation behaves similarly to *trans*-activation and, for Notch1, depends non-monotonically on ligand concentration (Figure 1E and Figure 2A,B). Together, these results strongly suggest that *cis*-activation is a core part of Notch signaling.

With the establishment of *cis*-activation, we next asked what developmental role *cis*-activation could play. Since Notch signaling promotes self-renewal and maintenance of stem cell fate in many systems (Imayoshi et al. 2010; Semerci et al. 2017), we chose to look at *cis*-activation in NSCs. We observed that survival of single isolated NSCs decreased significantly through inhibition of Notch signaling by DAPT treatment (Figure 3C), indicating that *cis*-activation can control NSC survival. Thus, self-renewing cells can be self-reliant, providing their own Notch signaling. This finding could help to explain how isolated stem cells can regenerate a complex tissue, as occurs in Lgr5+ intestinal crypt organoids (Sato et al. 2009) and mammary gland regeneration (Stingl et al. 2006), both of which are Notch-dependent.

Further expanding the functional possibilities for *cis*-activation, our modeling results suggest ways that *cis*-activation could widen the capabilities of the Notch pathway using only existing components. First, we showed how *cis*-activation could enable a “negative” Notch signaling modality in the presence of *trans*-ligand. At *cis*-Delta expression levels that produce high *cis*-activation, the introduction of an inhibitory *trans*-ligand led to strong, dose-dependent decreases in Notch activity (Figure 5K). This type of negative regulation is complementary to the *trans*-inhibition mechanism shown by (Benedito et al. 2009) where Notch1 activation by Dll4 was shown to be inhibited by the *trans*-ligand Jag1 during angiogenesis and in cell culture. Secondly, the model shows that using a combination of *cis*-activation, *cis*-inhibition, and *trans*-activation could in principle enable the cell to discriminate between the levels of its own (*cis*) ligands from those of its neighbors (*trans*) (Figure 5I). This property could be relevant for Notch-dependent fine-grained pattern formation through lateral inhibition circuits, in which cells coordinate their own Notch component levels with those of their neighbors (Collier et al. 1996; Barad et al. 2010; Sprinzak et al. 2011; Formosa-Jordan and Ibañes 2014b). More generally, the combination of *cis* and *trans* signaling can produce interesting behaviors. For example, in epidermal growth factor receptor (EGFR) signaling, autocrine and juxtacrine signaling modes lead to different biological outcomes (Raab and Klagsbrun 1997; Singh, Sugimoto, and Harris 2007). Similarly, in yeast, rewiring of the mating pathway to create an autocrine signaling system revealed that qualitatively different behaviors ranging from quorum sensing to bimodality could be generated by tuning the relative strengths of *cis* and *trans* signaling (Youk and Lim 2014). Looking forward, it will be interesting to see how Notch *cis*-activation and *trans*-inhibition mechanisms combine in natural developmental contexts.

Mechanistically, it remains puzzling how *cis* interactions could lead to both activation and inhibition in a ligand concentration-dependent fashion. Our study found that Notch ligand and receptor at the cell surface appears necessary both for productive *cis*-signaling and productive *trans*-signaling, and thus appears to be distinct from *cis*-interactions previously reported to occur within cellular endosomes (Coumailleau et al. 2009; Fürthauer and González-Gaitán 2009). Along with our observations that *cis*-and *trans*-signaling also share a similar dependence on ligand-receptor affinity and ligand concentrations, these results suggest that *cis*-activating complexes may resemble their *trans*-activating counterparts. Structural studies have shown that ligand-receptor binding could occur in both parallel and anti-parallel orientations (Cordle et al. 2008; Luca et al. 2015). It will therefore be interesting to see whether ligand-receptor complexes in different orientations can activate in a similar manner.

Mathematical modeling enabled us to explore different ways in which *cis*-activation and *cis*-inhibition could coexist. Critically, the simplest models in which the composition of activating and inhibitory complexes are identical cannot reproduce the shift from *cis*-activation to *cis*-inhibition (Figure 5A, Model 1). However, models in which the two complexes differ in their composition could more easily generate observed behaviors (Figure 5A, Model 2b-d)). These results suggest that *cis*-activation and *cis*-inhibition may involve the formation of distinct types of complexes. For example, one possibility is that *cis*-inhibition may involve the formation of larger ligand-receptor clusters than those involved in *cis*-activation.

The establishment of *cis*-activation as a prominent mode of signaling by the Notch pathway adds to our knowledge of how cells use Notch to communicate with itself and its neighbors, and suggests new ways in which cells can integrate different types of Notch signals. A more complete analysis of the *cis*- and *trans*-interactions among all ligand-receptor pairs, and for different levels of Fringe expression, could help to develop a more predictive understanding of how cells with distinct component combinations would be expected to activate both in *cis* and *trans*, thereby explaining how they use Notch to regulate their activities and those of their neighbors.

## Video Legends

### Video 1

#### Cis-activation of isolated engineered CHO-K1 cells

Examples of isolated CHO-K1 N1D1+Rfng cells activating prior to cell division in the *cis*-activation assay. (*Top row*) Blue channel shows fluorescence of the constitutively expressed nuclear H2B-Cerulean protein. (*Bottom row*) Green channel shows fluorescence of the Notch-activated H2B-Citrine reporter protein (also nuclear). The same intensity scales have been applied to each frame of the movie and for all cells. Interval between individual frames of the movie is 30 min. Not seen in the movie are non-fluorescent CHO-K1 cells, which surround each isolated fluorescent cell.

### Video 2

#### Cis-activation of isolated engineered NMuMG cells

Examples of isolated NMuMG N1D1+Rfng cells activating prior to cell division in the *cis*-activation assay. (*Top row*) Blue channel shows fluorescence of the constitutively expressed nuclear H2B-Cerulean protein. (*Bottom row*) Green channel shows fluorescence of the Notch-activated H2B-Citrine reporter protein (also nuclear). The fluorescence image is overlaid on the DIC image (grey), in which surrounding non-fluorescent NMuMG cells can be seen. The same intensity scales have been applied to each frame of the movie and for all cells. Interval between individual frames of the movie is 30 min.

## Source Data Legends

**Figure 5 – source data 1**

**Modeling code and data used in Figure 5.** Instructions are provided in the included README file.

## Materials and methods

### Plasmids

The majority of constructs used in this study have been previously described (Sprinzak et al. 2010). Briefly, the reporter for wild-type Notch activation was constructed from the 12xCSL plasmid (kind gift from U. Lendahl, Hansson et al. 2006), while the UAS reporter for Notch1ECD-Gal4 receptor activation was a kind gift from S. Fraser (de Celis, Bray, and Garcia-Bellido 1997). The construct containing the full-length human wild-type Notch1 sequence was a kind gift from J. Aster (de Celis and Bray 1997). The Notch1ECD-Gal4 plasmid was generated by replacing the Notch1ICD with amino acids 1–147 and 768–881 of the yeast Gal4. This construct was further modified by incorporating the sequence of the ankyrin (ANK) domain from Notch1ICD (amino acids 1872-2144) 3’ to the Gal4 sequence for use in the construction of the NMuMG cell lines. Design of the Notch2ECD-Gal4 plasmid was done in a manner similar to that of the Notch1ECD-Gal4 plasmid, but with incorporation of the expression cassette into a PiggyBac vector (System Biosciences, Palo Alto, CA) for efficient transfer to the cellular genome (Note: Notch1ECD-Gal4 was also incorporated into a PiggyBac vector when used in side-by-side comparisons with Notch2). The Notch ligand containing plasmids were based on the Tet-inducible system (Thermo Fisher Scientific, Waltham, MA). For the wild-type Notch1 cell line, we constructed a plasmid containing an inducible rat Dll1 coding sequence fused to the mCherry protein sequence. All other ligand plasmids were constructed containing an inducible human Dll1 or Dll4 coding sequence fused to a viral 2A sequence that allows for co-translation of a downstream H2B-mCherry protein sequence. In the NMuMG cells and in the CHO cell populations, the ligand plasmids were constructed within a PiggyBac vector. The Radical Fringe (Rfng) constructs used were based on those described by Lebon et. al. (LeBon et al. 2014). For use in CHO-K1 cells, Rfng was cloned into a pLenti expression construct from the ViraPower Lentiviral Expression System (Thermo Fisher Scientific), modified with a CMV promoter and a puromycin resistance gene. For use with the NMuMG cells, Rfng was cloned into an insulated UAS reporter construct (UAS surrounded by 2 copies of the 2xHS4 insulating element) by adding a separate cassette containing the sequence for the blastomycin resistance gene followed by a viral 2A sequence connected to the sequence for the rTetR gene fused to a HDAC4-2A-Rfng sequence. rTetR-HDAC4 (rTetS) was used to suppress constitutive Notch ligand expression in the NMuMG cells with the addition of Doxycycline to the cell media. The pCS-H2B-Cerulean plasmid was described in (Sprinzak et al. 2010). All cloning was done using standard molecular biology cloning techniques.

### Cell culture and transfections

#### CHO-K1

T-REx CHO-K1 cells from Thermo Fisher Scientific were cultured as described previously (Sprinzak et al. 2010; LeBon et al. 2014). Transfection of CHO-K1 cells was performed in 24-well plates with 800-1000 ng of DNA using the Lipofectamine LTX plasmid transfection reagent (ThermoFisher Scientific). 24 hours post-transfection, cells were split into new 6-well plates and cultured for 1-2 weeks in media containing 400 ug/ml Zeocin, 600 ug/ml Geneticin, 300 ug/ml Hygromycin, 10 ug/ml Blasticidin, or 3 ug/ml Puromycin as appropriate, and surviving transfected cells were either used as clonal populations or subcloned by limiting dilution.

#### NMuMG

NMuMG cells (ATCC, Manassas, VA) were cultured using the manufacturer’s recommended culturing protocol with the addition of 1 mM Sodium Pyruvate and 100 U ml-1 penicillin, 100 μg ml-1 streptomycin (Thermo Fisher Scientific) to the media. Transfection, selection and clonal isolation of NMuMG cells was performed similarly to CHO-K1 cells.

#### Caco-2

Caco-2 C2BBe1 cells from ATCC were cultured in Dulbecco’s modified Eagle’s medium (DMEM, Thermo Fisher Scientific) supplemented with 10% Tet-System Approved Fetal Bovine Serum (Takara Bio USA Inc., Mountain View, CA), 2 mM L-Glutamine, 100 U ml-1 penicillin, 100 μg ml-1 streptomycin, 1 mM sodium pyruvate, and 1X MEM Non-Essential Amino Acids Solution (Thermo Fisher Scientific). Transfections were performed by following the Thermo Fisher Lipofectamine LTX protocol optimized for Caco-2 cells. 24 hours after transfection, Caco-2 cell populations were plated for experiments.

#### Neural Stem Cells

Neural stem cells derived from the E14.5 mouse cortex were purchased from EMD Millipore (Burlington, MA, Catalog No. SCR029) and cultured according to the manufacturer’s protocols. Briefly, tissue-culture surfaces were coated overnight with poly-L-ornithine (10 ug/ml, Sigma Aldrich Catalog No. P3655) and Laminin (Sigma Aldrich Catalog No. L2020). For standard culture, cells were then plated in Neurobasal medium (EMD Millipore, Catalog No. SCM033) in the presence of 20 ng/ml recombinant FGF (EMD Millipore Catalog No. GF003), 20 ng/ml EGF (Millipore Catalog No. GF001), and Heparin (Sigma Catalog No. H3149). Cells were passaged using ESGRO Complete Accutase (Millipore Catalog No. SF006), cryo-preserved in medium + 10% DMSO, and typically used for experiments within six passages.

### Lentiviral production and infection

Lentivirus was produced using the ViraPower Lentiviral Expression System (Thermo Fisher Scientific). Briefly, 293FT producer cells were transfected with our pLenti expression construct along with the packaging plasmid mix. 48 hours post-transfection, virus containing cell media was collected, centrifuged to remove cell debris and filtered through a 0.45 um filter (EMD Millipore). Viral supernatant was added 1:2 to sparsely plated CHO-K1 cells in a total volume of 400 ul media in a 24-well plate and incubated at 37°C, 5% CO_2_. 24 hours post-infection, virus-containing media was removed, and cells were plated under limiting dilution conditions in 96-well plates for clonal selection. Expression of the integrated gene was checked by qRT-PCR analysis.

### CRISPR-Cas9 knock-out of endogenous NMuMG Notch2 and Jagged1

Endogenous Notch2 and Jagged1 genes were knocked out in NMuMG cells using the CRISPR-Cas9 plasmid system developed by the Zhang Lab at MIT (Cong et al. 2013). Cloning was done according to the published protocol using the pX330 plasmid and inserting a guide sequence using the following oligos for targeting mouse Notch2 or Jagged1:

Notch2

mN2 C2 OligoF: 5’-CACCGGGTGGTACTTGTGTGCCGCA-3’
mN2 C2 OligoR: 5’-AAACTGCGGCACACAAGTACCACCC-3’

Jagged1

mJ1 C1 OligoF: 5’-CACCGCGGGTGCACTTGCGGTCGCC-3’
mJ1 C1 OligoR: 5’-AAACGGCGACCGCAAGTGCACCCGC-3’

The guide sequence modified pX330 plasmids were transfected into NMuMG cells using the standard Lipofectamine LTX protocol (Thermo Fisher Scientific). 48 hours post-transfection, genomic DNA was harvested from the cell population, and guide sequence function was analyzed using the SURVEYOR Mutation Detection Kit (Integrated DNA Technologies Inc., Skokie, IL). After genomic knock-out mutation was verified, transfected cells were placed under clonal selection using limiting dilution. Genomic DNA was isolated from clones and used to PCR amplify targeted sequences using the following primers (Integrated DNA Technologies Inc.):

<u>Notch2</u>

mN2 C2F: 5’-GTCACCCGTCTGGTATTTTGTTAC-3’
mN2 C2R: 5’-GAGCTGCTGTGATCGAAGTG-3’

<u>Jagged1</u>

mJ1 C1F: 5’-CCAAAGCCTCTCAACTTAGTGC-3’
mJ1 C1R: 5’-CTTAGTTTTCCCGCACTTGTGTTT-3’

PCR products were purified using the QIAquick PCR Purification Kit (Qiagen, Hilden, Germany) and sent for sequencing (Laragen Inc., Culver City, CA) to determine clones that contained gene knock-out mutations. Gene knock-out was also verified by staining mutated cells with antibodies against Notch2 and Jagged1 (anti-mouse Notch2-PE and anti-mouse Jagged1-PE, BioLegend, San Diego, CA) in order to determine their levels of surface expression. Cells were analyzed by flow cytometry (see flow cytometry section below).

### Availability assay for Notch1ECD-Gal4 or Notch1ECD-Gal4-ANK in NMuMG cells

Surface staining of either Notch1ECD-Gal4 or Notch1ECD-Gal4-ANK was performed using the availability assay as described previously (LeBon et al. 2014). Briefly, cells were washed in phosphate buffered saline (PBS) and blocked in a PBS solution containing 2% BSA and 100 ug/ml CaCl_2_ while rocking for 40 minutes at room temperature. After blocking, cells were rocked in a PBS solution containing 2% FBS, 100 ug/ml CaCl_2_, and 10 ug/ml of recombinant mouse Dll1-Fc (rmDll1-Fc) for 1 hour at room temperature. The recombinant Dll1 protein can bind to the available (free) Notch receptors at the cellular surface. After the 1 hour incubation, cells were washed 3x with PBS and incubated with a secondary antibody conjugated to an Alexa Fluor 488 dye. Cells were rocked at room temperature for 1 hour, washed 3x with PBS, and Notch localization on the cell surface was imaged on an EVOS FL Auto Cell Imaging system (Thermo Fisher Scientific).

### qRT-PCR

Expression of Radical Fringe (Rfng) in clones was determined by quantitative RT-PCR. RNA was isolated from clonal cells using the RNeasy Mini Kit (Qiagen) following the manufacturer’s protocol. 200-500 ng RNA was used to make cDNA using the iScript cDNA Synthesis Kit (Bio-Rad Laboratories, Hercules, CA). 2 ul of cDNA was used in the qPCR reaction along with SsoAdvanced Universal Probes Supermix (Bio-Rad Laboratories) and primers/probes (Integrated DNA Technologies Inc.) as follows:

<u>Rfng</u>

Probe: 5′-6-FAM/ZEN/3’ IB®FQCTCGTGAGATCCAGGTACGCAGC-3’
Primer 1: 5′-TCATTGCAGTCAAGACCACTC-3’
Primer 2: 5′-CGGTGAAAATGAACGTCTGC-3’

<u>b-Actin</u> (Housekeeping Gene for CHO-K1 cells)

Probe: 5′-HEX /ZEN/ 3’ IB®FQ-ACCACACCTTCTACAACGAGCTGC-3’
Primer1: 5′-ACTGGGACGATATGGAGAAG-3’ Primer2: 5′-GGTCATCTTTTCACGGTTGG-3’

<u>GAPDH</u> (Housekeeping Gene for NMuMG cells)

Probe: 5′-HEX /ZEN/ 3’ IB®FQ-AGGAGCGAGACCCCACTAACATCA-3’
Primer1: 5’-CTCCACGACATACTCAGCAC-3’
Primer2: 5’-CCACTCACGGCAAATTCAAC-3’

Samples were run in triplicate on a CFX96 Touch Real-Time PCR Detection System (Bio-Rad Laboratories), and relative gene expression levels were calculated using the standard delta-delta Cq method.

### Plate-bound Dll1

Coating of tissue culture plates with recombinant human Dll1^ext^-Fc fusion proteins (kind gift from I. Bernstein) was done as previously reported (Nandagopal et al. 2018). Briefly, Dll1^ext^-Fc proteins were diluted to 2.5 ug/ml in 1xPBS (Thermo Fisher Scientific), and the solution was used to coat the tissue-culture surface for 1 hour at room temperature with rocking. Post-incubation, the solution was removed and cells were plated for the experiment.

### RNA sequencing of NSCs

NSCs were cultured for 12h in high (20 ng/ml EGF, 20 ng/ml FGF) or low (0.5 ng/ml EGF) growth factor with or without 10 μM DAPT. RNA was extracted using the RNeasy kit (QIAGEN) and submitted to the Caltech sequencing core facility, where cDNA libraries for RNaseq were prepared according to standard Illumina protocols. 50 base single-end read (50SR) sequencing was performed on a HiSeq2500 machine at the same facility. Reads were assembled, aligned, and mapped to the mouse genome (mm10 assembly) on the Galaxy server, using RNA Star. Cufflinks was used to calculate FPKM values. Raw and processed sequencing data is available in GEO (Accession GSE113937).

### Single molecule HCR-FISH detection of Notch targets in NSCs

#### Experimental Protocols

NSCs, cultured in standard high growth factor containing media (see cell culture and transfections section above), were plated on 10 ug/ml poly-L-ornithine and 50 ug/ml Laminin-coated glass plates, at a surface density of ~4 cells/mm2. At the time of plating, cells were transferred to low growth factor conditions (0.1 ng/ml EGF, 5 ug/ml Heparin, no FGF), with or without 10 μM DAPT. 6 hours post plating, cells were fixed using 4% formaldehyde. Prescribed protocols were followed for hybridizing DNA probes to targets genes (*Hes1*, *Hey1*, and *Hes5*) and amplifying bound probes (Choi et al. 2018). Briefly, fixed cells were incubated overnight at 37°C with 10 pairs of probes per gene diluted in 30% formamide-containing buffer. Subsequently, probes were removed and cells were washed at 37°C. Then, DNA amplifiers, designed to detect bound probes and coupled to one of three Alexa Fluor dyes (488, 546, or 647), were added to the sample, and amplification allowed to proceed at room temperature for ~50 min. Samples were then washed in high salt solution (5x SSC with Tween), and stained with DAPI, prior to imaging.

#### Imaging

Samples were imaged at 60x (1.3 NA, oil) on an inverted epi-fluorescence microscope (Nikon Ti: Nikon Instruments Inc., Melville, NY) equipped with an LED lightsource (Lumencor, Beaverton, OR) and hardware autofocus. Fields of view that contained between 1-3 well-separated cells were picked manually and Z-stacks were acquired over 16 μm at each position.

#### Analysis

Custom MATLAB (2015a, Mathworks) software was used to semi-automatically segment cells based on autofluorescence in the 488 channel. mRNA transcripts typically appeared as 3-5 voxel-wide high-intensity dots in the images. Previously used MATLAB software for detecting dots (Frieda et al. 2017) was adapted to automatically detect dots in images based on user-defined thresholds. For direct comparison, the same thresholds were applied to data from DAPT-treated and untreated samples.

### Cis-activation and relative density assays

CHO-K1 engineered cell lines were pre-incubated with 1uM of the gamma-secretase inhibitor DAPT (Sigma-Aldrich, St. Louis, MO) and various concentrations of the tetracycline analog, 4-epiTetracycline (4-epiTc, Sigma-Aldrich) 48 hours prior to the setup of assays. For the *cis*-activation assay, cells were washed to remove DAPT, counted, and plated sparsely at 5K cells per 24-well plate, surrounded by 150K wild-type CHO-K1 cells. 4-epiTc was added back into the media (0, 20, 35, 50, 80, or 200 ng/ml) and the cells were either incubated at 37°C, 5% CO_2_ for <24 hours before analysis by flow cytometry or imaged using time-lapse microscopy. For the ‘control’ *cis*-activation assay, 4K 4-epiTc pre-induced N1D1+Rfng cells were plated along with 4K N1 receiver cells (no ligand present), and 750K CHO-K1 cells per 6-well plate. Cells were analyzed for activation by flow cytometry as previously mentioned. Relative density assays were performed using the same setup conditions as the *cis*-activation assay, but with varying ratios of engineered:wild-type cells plated. Keeping total cell numbers at 150K cells per 24-well plate, either 5K, 10K, 25K, 50K, 75K or 100K engineered cells were plated along with wild-type CHO-K1 cells. For the *cis*-activation and relative density assays using Blebbistatin treated cells, assay setup was done exactly as mentioned above but with the addition of 10 uM (±)-Blebbistatin (Sigma-Aldrich) added to the cell media at the time of plating.

For NMuMG cells, cis-activation and density assays were performed just like those with CHO-K1 cells. However, cells were pre-incubated in 10 uM DAPT with or without the addition of 1 ug/ml or 10 ug/ml Doxycycline (Takara Bio USA Inc.) for 3 days in order to decrease ligand expression levels prior to assay setup. Cells were plated with or without Dox, with the addition of 100 ng/ml Dexamethasone (Sigma-Aldrich) for each assay.

Caco-2 cells were pre-incubated with 100 uM DAPT for 1 day prior to transfection. 24 hours post-12xCSL reporter transfection, the cells were washed, counted and plated sparsely at 3.5K cells in a 24-well plate for the *cis*-activation assay or 7-fold more dense at 3.5K cells in a 96-well plate for a density assay with or without the addition of DAPT. <24 hours after plating, cells were analyzed by FLOW cytometry.

### Cis-activation assay in suspension

For performing the *cis*-activation assay with cells in suspension, 24-well 10mm diameter glass No. 1.5 coverslip plates (MatTek Corp., Ashland, MA) were coated with the siliconizing reagent Sigmacote (Sigma) to prevent cells from adhering to the plate surface. Cells were plated as mentioned previously and placed on a rocker at 37oC, 5% CO2 overnight before analysis by flow cytometry.

### Notch receptor/ligand blocking assay

Engineered CHO-K1 cells were pre-incubated in 1 uM DAPT, with and without the addition of 4-epiTc, for 2 days prior to the start of the assay. Cells were then incubated with either 10 ug/ml mouse IgG_2A_ control protein or 10 ug/ml mouse Notch1 Fc chimera protein (R&D Systems, Minneapolis, MN) along with DAPT and 4-epiTc overnight at 37°C, 5% CO_2_. The next day, cells were washed, counted, and plated at 5K cells per 24-well plate along with 150K wild-type CHO-K1 cells for a *cis*-activation assay, or at 150K cells per 24-well plate for a relative density assay with the addition of 4-epiTc. Cells grown similarly, but in the absence of 4-epiTc, were used as a control. <24 hours post-plating, cells were analyzed by flow cytometry.

### NSC survival assay

#### Experimental Protocols

NSCs, cultured in standard high growth factor containing media (see cell culture and transfections section above), were plated on 10 ug/ml poly-L-ornithine and 5 ug/ml Laminin-coated plastic surfaces (12-well TC-treated plates, Corning Inc.) at a surface density of ~20 cells/mm^2^. At the time of plating, cells were transferred to low growth factor conditions (0.1 ng/ml EGF, 5 ug/ml Heparin, no FGF), with or without 10 μM DAPT. 12 hours post plating, cells were fixed using 4% formaldehyde. Samples were then stained with DAPI prior to imaging.

#### Imaging

Samples were imaged at 20x (0.75 NA, air) in an inverted epi-fluorescence microscope (Olympus IX81) equipped with an LED lightsource (XCite LED) and hardware ZCD2 autofocus. 484 600 μm × 600 μm fields of view were acquired from across the well for each sample.

#### Analysis

Custom MATLAB (2015a, Mathworks) software was used to automatically segment nuclei based on DAPI staining. The number of nuclei were then counted for each of the different samples.

### NSC differentiation assay

#### Experimental Protocols

NSCs, cultured in standard high growth factor containing media (see cell culture and transfections section above), were plated on 10 ug/ml poly-L-ornithine and 5 ug/ml Laminin-coated plastic surfaces (6-well TC-treated plates, Corning Inc.) at a surface density of ~20 cells/mm2. At the time of plating, cells were transferred to differentiation conditions (1% FBS + 1% N2 supplement + 2% B27 supplement + 1 mM GlutaMax supplements; supplements purchased from Thermo Fisher), with or without 10 μM DAPT. 24 hours post plating, cells were fixed using 4% formaldehyde. Samples were blocked in 2% BSA solution + 0.3% Triton-X, then incubated overnight at 4°C with 1:500 mouse anti-GFAP (GA5, Catalog #3760, Cell Signaling Technology, Danver, MA) and 1:1000 rabbit anti-β-III-Tubulin (Catalog No. ab18207, Abcam, Cambridge). Cells were subsequently washed and incubated with DAPI and anti-mouse or -rabbit 1:1000 secondary antibodies conjugated to AlexFluor 488 and 594, respectively, in blocking solution for 1 hour at room temperature, before imaging.

#### Imaging

Samples were imaged at 20x (0.75 NA, air) in an inverted epi-fluorescence microscope (Olympus IX81) equipped with an LED lightsource (XCite LED) and hardware ZDC2 autofocus.

#### Analysis

Custom MATLAB (2015a, Mathworks) software was used to automatically segment cells based on DAPI staining and fluorescence in the 594 (β-III-Tubulin) channel, which shows detectable cell-wide staining in all cells. Total fluorescence in the 488 and 594 channels were calculated in each cell segment.

### Time-lapse setup, image acquisition and analysis

#### Experimental setup

For imaging, CHO-K1 cells were plated in 24-well 10mm diameter glass No. 1.5 coverslip plates (MatTek Corp.) coated with 5 ug/ml hamster Fibronectin (Oxford Biomedical Research, Rochester Hills, MI) in complete cell media. NMuMG cells were plated in 24-well tissue culture treated ultrathin glass film bottom plates (Eppendorf, Hamburg, Germany) in complete cell media.

#### Acquisition

Movies were acquired at 20X (0.75 NA) on an Olympus IX81 inverted epifluorescence microscope (Olympus, Tokyo, Japan) equipped with hardware autofocus (ZDC2), an iKon-M CCD camera (Andor, Concord, MA) and an environmental chamber maintaining cells at 37°C, 5% CO_2_ with humidity throughout the length of the movie. Automated acquisition software (Metamorph, Molecular Devices, San Jose, CA) was used to acquire images every 30 min in multiple channels (YFP, RFP, CFP) or differential interference contrast (DIC), from multiple stage positions.

#### Analysis

Custom MATLAB code (2013a, MathWorks) was used to segment cell nuclei in images based on constitutive Cerulean fluorescence. Briefly, the segmentation procedure uses built-in edge detection MATLAB functions and adaptive thresholds to detect nuclear segments. Nuclear segments were then matched in pairs of images corresponding to consecutive time frames, and thus tracked through the duration of the movie. Single-cell tracks were subsequently curated manually to correct for errors in segmentation/tracking. Fluorescence data was extracted from nuclear segments by calculating the integrated fluorescence within the segment and subtracting a background fluorescence level estimated from the local neighborhood of the segment. This fluorescence was linearly interpolated across time frames where nuclei could not be segmented automatically. Division events were detected automatically, and fluorescence traces were corrected for cell division by adding back fluorescence lost to sister cells. The resulting ‘continuized’ traces were smoothed and the difference in fluorescence between consecutive time frames was calculated. A smoothed version of this difference was used as an estimate of production rate of the fluorescent protein.

### Flow cytometry analysis

For analysis of cells by flow cytometry, cells were trypsinized in 0.25% Trypsin-EDTA (Thermo Fisher Scientific) and resuspend in 1x Hanks Balanced Salt Solution (Thermo Fisher Scientific) supplemented with 2.5 mg/ml Bovine Serum Albumin (Sigma-Aldrich). Resuspended cells were filtered using 40 um cell strainers (Corning Inc., Corning, NY) into U-bottom 96-well tissue-culture treated plates. Cells were analyzed on a MACSQuant VYB Flow Cytometer (Miltenyi Biotech, <u>Bergisch Gladbach, Germany</u>) located at the Caltech Flow Cytometry Facility (Caltech, Pasadena, CA). Data was analyzed in MATLAB using custom software (EasyFlow) (Antebi et al. 2017), and forward and side-scatter profiles were used to gate on the proper cell populations. Fluorescence intensity of single-cells was measured for each appropriate channel.

### Mathematical models – see also Figure 5 – source data 1

#### Models

The models analyzed here attempt to recapitulate the behavior of the system at steady state. Components of the system include free Notch receptor (N) and free Delta ligand (D) and, depending on the model, cis- and trans complexes (C^+^/C^−^, and T, respectively) between ligands and receptors. In all models, N and D are produced at a rate of α_N_ and α_D_, and degraded at the rate of γ_N_ and γ_D_, respectively.

I) **Model 0**. This model assumes that N and D interact at a rate *k*^+^ to produce a single type of cis-complex, C^+^, which dissociates at a rate *k^−^*, and is degraded at a rate γ_C+_. That is,

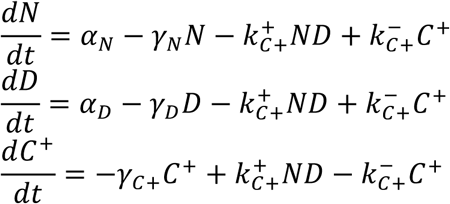
II) **Model 1**. This model assumes that N and D can interact to produce two distinct *cis*-complexes, active C^+^ and inactive C^−^. These complexes are formed through similar second-order interactions between N and D, occurring with different rate coefficients. They similarly dissociate and degrade at different rates, and cannot interconvert.

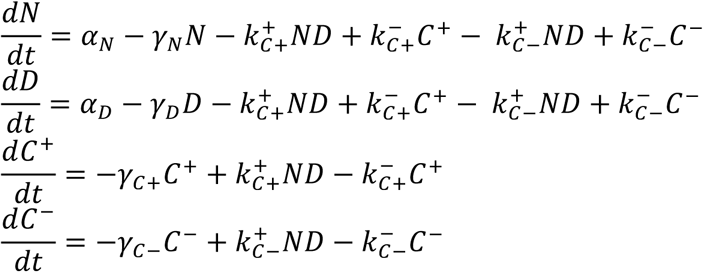
III) **Models 2a-d**. These models also assume that N and D can interact to produce two distinct *cis*-complexes, C^+^ and C^−^. However, in these models, C^−^ requires C^+^ for its formation (a-c) or the stoichiometry of inactive C^−^ formation is higher than that of C^+^ (d).

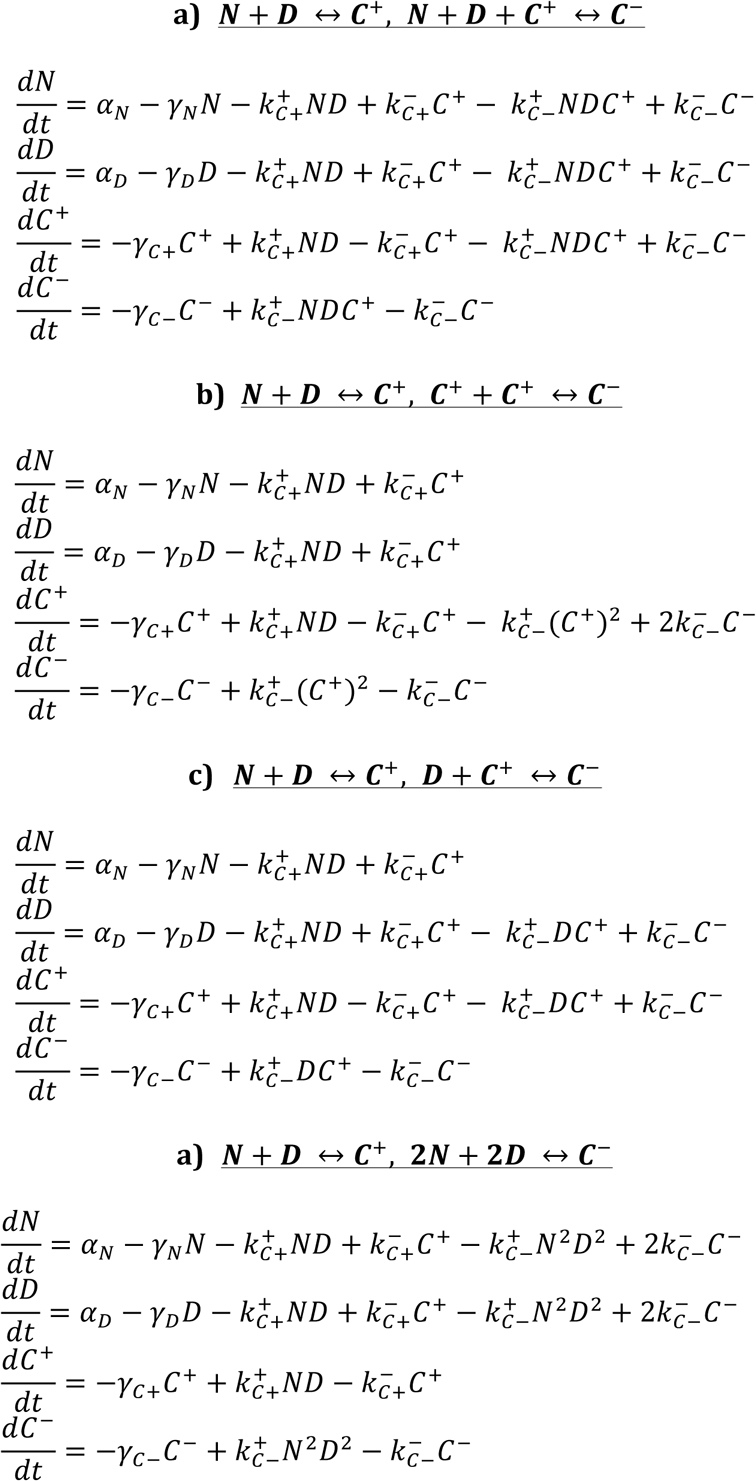
IV) **Model 2c + trans-interactions**: To model the effect of trans-interactions in the context of model 2c, it was assumed that the *trans*-complex T is formed through interactions between N and *trans* Delta (D_*trans*_, assumed to be constant). The rate coefficients of its formation, dissociation, and degradation are assumed to be the same as that of C^+^.

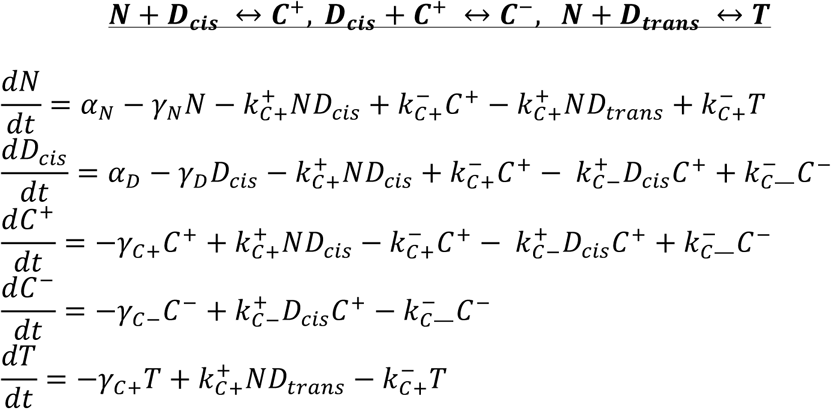

#### Parameter scan, numerical simulations, and analyses

Model 0 contains 6 parameters (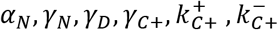), while Models 1 and 2 contain 3 additional parameters (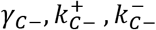). Using the built-in *lhsdesign* function in MATLAB (2015a, Mathworks), the Latin Hypercube Sampling algorithm was applied to pick 10,000 parameters, each in the range 10^−2^ to 10^2^. For each parameter set, the model was simulated for each of 10 values of *α_D_*, logarithmically spanning a 100-fold range around the sampled value of α_*N*_. The *fsolve* function, with initial conditions [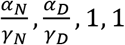] for N, D, C^+^, C^−^, respectively, was used to numerically approximate the steady state solution for each parameter set.

For each solution, the following features of the *α_D_* vs. C^+^ profile were calculated: the relative value of *α_D_* at which C^+^ was maximum (‘C-max’), and the fractional increases in C-max relative to its value at the lowest and highest values of *α_D_*. Parameters that produced C^+^ profiles that peaked between the 1^st^ and 8^th^ value of *α_D_* were deemed to be non-monotonic.

For the *trans*-interaction model, first the values of D_cis_ obtained at D_*trans*_ = 0 were calculated for each parameter set. For subsequent simulations, the values of D_*trans*_ were chosen to be the same as that of these D_*cis*_ values, i.e., D_*cis*_ produced in the absence of *trans*-ligand.

### Statistics

No statistical method was used to determine sample sizes. The sample sizes used were based on general standards accepted by the field. The number of replicates used for each experimental analysis is listed in the figure legends. All replicates are biological replicates, corresponding to measurements performed on distinct biological samples, as opposed to performing the same tests multiple times on a single sample (technical replicates). *P*-values for Figure 1 and 3 were calculated using the two-sided KS-test. All pairwise comparisons between samples fulfilled the criterion n1*n2/(n1 + n2) ≥ 4, where n1 and n2 represent the number of data points in two samples. Under this condition, the KS-statistic is greater than twice the inverse of the Kolmogorov statistic, and the calculated *P*-value is accurate. The non-parametric nature of the KS-test obviates the need to make assumptions regarding the shape of the distributions being compared. *P*-values for Caco-2 cell measurements (Figure 1-figure supplement 4) and CHO cell surface binding assay measurements (Figure 4) were calculated using the one-sided Student T-test, which assumes that random error in the measurement follows a normal (Gaussian) distribution.

## Acknowledgements

This work was supported by Howard Hughes Medical Institute (M.B.E.), the Defense Advanced Research Projects Agency under Contract No. HR0011-16-0138, the National Institutes of Health grant R01 HD075335 and the NSF under grant EFRI 1137269. N.N was a Howard Hughes Medical Institute International Student Research fellow. We thank Pulin Li, Mark Budde, Heidi Klumpe, Ronghui Zhu, Rachael Kuintzle, Laurent Potvin-Trottier and James Linton for critical feedback on the manuscript. Harry Choi and Colby Calvert, Caltech Flow Cytometry Facility, Caltech Biological Imaging Facility, and the Millard and Muriel Jacobs Genetics and Genomics Laboratory at Caltech provided essential technical assistance.

## Supplementary materials

**Supplementary File 1A.** Cell lines used in this work.

| **Cell line name** | **Selection** | **Notes** | **Figure(s)** |
| --- | --- | --- | --- |
| CHO-K1 | Blasticidin 10µg/mL | Base wild-type cell line expressing TetR (K1-TREx CHO-K1) | **1,2,4**<br>**1-figure supplement 1,2,3**<br>**2-figure supplement 1,2**<br>**4-figure supplement 1** |
| N1D1 | Blasticidin 10µg/mL,<br>Zeocin 400µg/mL,<br>Geneticin 600µg/mL<br>Hygromycin 500µg/mL | CHO-K1 expressing pEF-hNECD-Gal4esn + pcDNA5-TO-DII1-T2A-H2B-mCherry + pEV-UAS-H2B-Citrine | **2** |
| N1D1+Rfng | Blasticidin 10µg/mL,<br>Zeocin 400µg/mL,<br>Geneticin 600µg/mL,<br>Hygromycin 500µg/mL,<br>Puromycin 3µg/mL | CHO-K1 expressing pEF-hNECD-Gal4esn + pcDNA5-TO-DII1-T2A-H2B-mCherry + pEV-UAS-H2B-Citrine + pLenti-CMV-R-fringe-T2A-Puromycin | **1,2,4**<br>**1-figure supplement 1,2**<br>**4-figure supplement 1** |
| N1<sup>WT</sup>D1+Rfng | Blasticidin 10µg/mL,<br>Zeocin 400µg/mL,<br>Geneticin 600µg/mL,<br>Hygromycin 500µg/mL,<br>Puromycin 3µg/mL | CHO-K1 expressing pcDNA3-hN1-mod1 + pcDNA5-TO-DII1-mCherry + pEV-12xCSL-H2B-Citrine + piggyBac-CMV-R-fringe + pCS-H2B-Cerulean | **1-figure supplement 3** |
| NMuMG<br>N1D1+Rfng | Blasticidin 10µg/mL,<br>Zeocin 400µg/mL,<br>Geneticin 600µg/mL, | NMuMG expressing piggyBac- CMV-hNECD-Gal4-ANK-T2A-H2B- | **1-figure supplement 3** |
|  | Hygromycin<br>300µg/mL, 4-epiTc<br>200ng/ml | Cerulean + piggyBac-CMV-TO Dll1-T2A-H2B-mCherry-P2A-Hygromycin + pEV-2xHS4-UAS-H2B-Citrine-T2A-tTS-2xHS4-Blast-T2A-rTetR-HDAC4-P2A-R-fringe |  |
| Caco-2 | None | Wild-type cells used to<br>transfect in pEV-12xCSL-H2B-Citrine | **1-figure**<br>**supplement 4** |
| N1D1 Pop | Blasticidin 10µg/mL,<br>Zeocin 400µg/mL,<br>Geneticin 600µg/mL,<br>Hygromycin 300µg/mL | CHO-K1 with pEV-UAS-H2B-Citrine + pCS-H2B-Cerulean<br>+ piggyBac-TO-Dll1-T2A-H2B-mCherry + piggyBac-CMV-hN1ECD-Gal4<br>Cell population (Pop) | **2** |
| N1D4 Pop | Blasticidin 10µg/mL,<br>Zeocin 400µg/mL,<br>Geneticin 600µg/mL,<br>Hygromycin 300µg/mL | CHO-K1 with pEV-UAS-H2B-Citrine + pCS-H2B-Cerulean<br>+ piggyBac-TO-Dll4-T2A-H2B-mCherry + piggyBac-CMV-hN1ECD-Gal4<br>Cell population (Pop) | **2**<br>**2-figure**<br>**supplement 1** |
| N2D1 Pop | Blasticidin 10µg/mL,<br>Zeocin 400µg/mL,<br>Geneticin 600µg/mL,<br>Hygromycin 300µg/mL | CHO-K1 with pEV-UAS-H2B-Citrine + pCS-H2B-Cerulean<br>+ piggyBac-TO-Dll1-T2A-H2B-mCherry + piggyBac-CMV-hN2ECD-Gal4<br>Cell population (Pop) | **2**<br>**2-figure**<br>**supplement 2** |
| N2D4 Pop | Blasticidin 10µg/mL,<br>Zeocin 400µg/mL,<br>Geneticin 600µg/mL,<br>Hygromycin 300µg/mL | CHO-K1 with pEV-UAS-H2B-Citrine + pCS-H2B-Cerulean<br>+ piggyBac-TO-Dll4-T2A-H2B-mCherry + piggyBac-CMV-hN2ECD-Gal4<br>Cell population (Pop) | **2**<br>**2-figure**<br>**supplement 2** |
| NSC | None | E14.5 mouse neural cortical<br>stem cells (NSC) | **3**<br>**3-figure**<br>**supplement**<br>**1,2** |

**Supplementary File 1B.** Plasmids used in this work

| Construct name | Promoter | Gene | Mammalian selection | Role |
| --- | --- | --- | --- | --- |
| pEV-UAS-H2B-Citrine | UAS | H2B-Citrine | Zeocin | Reporter for Notch1ECD-Gal4 receptor in CHO cells |
| pEV-2xHS4-UAS-H2B-Citrine-T2A-tTS-2xHS4-Blast-T2A-rTetR-HDAC4-P2A-R-fringe | UAS for H2B-Citrine-T2A-tTS<br>SV40 for Blast-T2A-rTetR-HDAC4-P2A-R-fringe | 2xHS4 insulator elements surrounding H2B-Citrine with co-translationally expressed tTS<br>Blasticidin with co-translationally expressed rTetR-HDAC4 and R-fringe | Blasticidin | Reporter for Notch1ECD-Gal4-ANK receptor in NMuMG cells (tTS was not relevant for this work and was inactivated by 4-epiTc)<br>rTetR-HDAC4 was used to decrease Delta expression in the presence of Dox.<br>Constitutive R-fringe expression in NMuMG cells |
| pEV-12xCSL-H2B-Citrine | 12xCSL | H2B-Citrine | Zeocin | Reporter for Notch1<sup>WT</sup> receptor in CHO and Caco-2 cells |
| pEF-hN1ECD-Gal4 | pEF | hNotch1ECD-Gal4 | Neomycin/Geneticin | Notch1ECD-Gal4 synthetic receptor used in CHO clones |
| piggyBac-CMV-hN1ECD-Gal4 | CMV | hNotch1ECD-Gal4 | Neomycin/Geneticin | Notch1ECD-Gal4 synthetic receptor used in CHO populations |
| piggyBac-CMV-hN2ECD-Gal4 | CMV | hNotch2ECD-Gal4 | Neomycin/Geneticin | Notch2ECD-Gal4 synthetic receptor used in CHO populations |
| pcDNA3-hN1-mod1 | CMV | hNotch1 (full-length) | Neomycin/Geneticin | Wild-type Notch1 receptor used in CHO clones |
| piggyBac-CMV-hNECD-Gal4-ANK-T2A-H2B- | CMV | hNotch1ECD-Gal4-ANK (Notch1ICD Ankyrin domain) with a co-translationally | Neomycin/Geneticin | Notch1ECD-Gal4-ANK synthetic receptor used in NMuMG clones |
| Cerulean |  | expressed H2B-Cerulean |  |  |
| pcDNA5-TO-DII1-T2A-H2B-mCherry | CMV-TO | Delta-like1 with a co-translationally expressed H2B-mCherry | Hygromycin | Inducible Delta-like1 ligand used in CHO clones and populations |
| pcDNA5-TO-DII1- mCherry | CMV-TO | Delta-like1 fused to mCherry | Hygromycin | Inducible Delta-like1-mCherry fusion used in CHO clones with Notch1<sup>WT</sup> receptor |
| piggyBac-CMV-TO DII1-T2A-H2B-mCherry-P2A-Hygromycin | CMV-TO | Delta-like1 with a co-translationally expressed H2B-mCherry and hygromycin resistance gene | Hygromycin | Inducible Delta-like1 ligand used in CHO populations and NMuMG clones |
| piggyBac-CMV-TO-DII4-T2A-H2B-mCherry-P2A-Hygromycin | CMV-TO | Delta-like4 with a co-translationally expressed H2B-mCherry and hygromycin resistance gene | Hygromycin | Inducible Delta-like4 ligand used in CHO populations |
| pLenti-CMV-R-fringe-T2A-Puromycin | CMV | R-fringe with a co-translationally expressed puromycin resistance gene | Puromycin | Constitutive R-fringe expression in CHO N1D1+Rfng cells |
| pCS-H2B-Cerulean | CMV | H2B-Cerulean | None | Segmentation color used in CHO cells |

**Figure 1-figure supplement 1.**
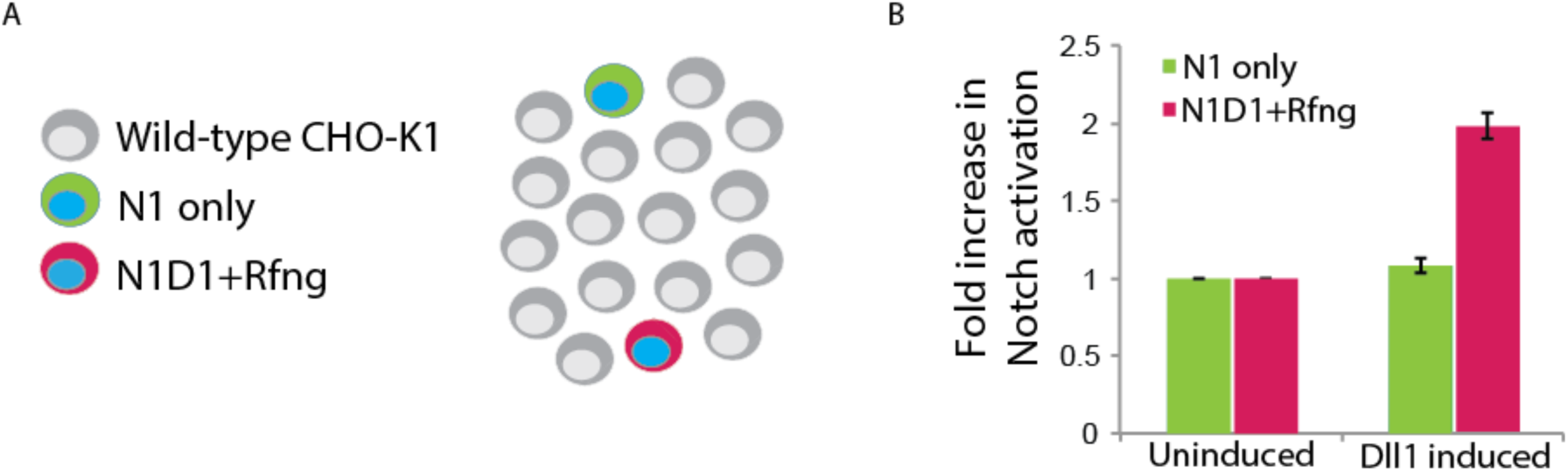
Cis-activation assay enables isolation of individual engineered cells. **(A)** Schematic of ‘control’ *cis*-activation assay used to verify that the relative density of cells was low enough to prevent *trans*-interactions. N1D1+Rfng cells (magenta, 0.5%) were mixed with Notch receiver cells (‘N1 only’, green, 0.5%), and plated with an excess of wild-type CHO-K1 cells (grey), with or without addition of 80 ng/ml 4-epiTc for Dll1 induction. Blue nuclei represent constitutive H2B-Cerulean expression. **(B)** Mean activation levels (measured by flow cytometry) in N1D1+Rfng (magenta) and N1-only (green) cells. Data was normalized to background Citrine levels in N1D1+Rfng cells not induced to express Dll1 (‘uninduced’). Dll1 expression increases activation of N1D1+Rfng cells, but not N1-only cells. Error bars represent the s.e.m across n=3 biological replicates.

**Figure 1-figure supplement 2.**
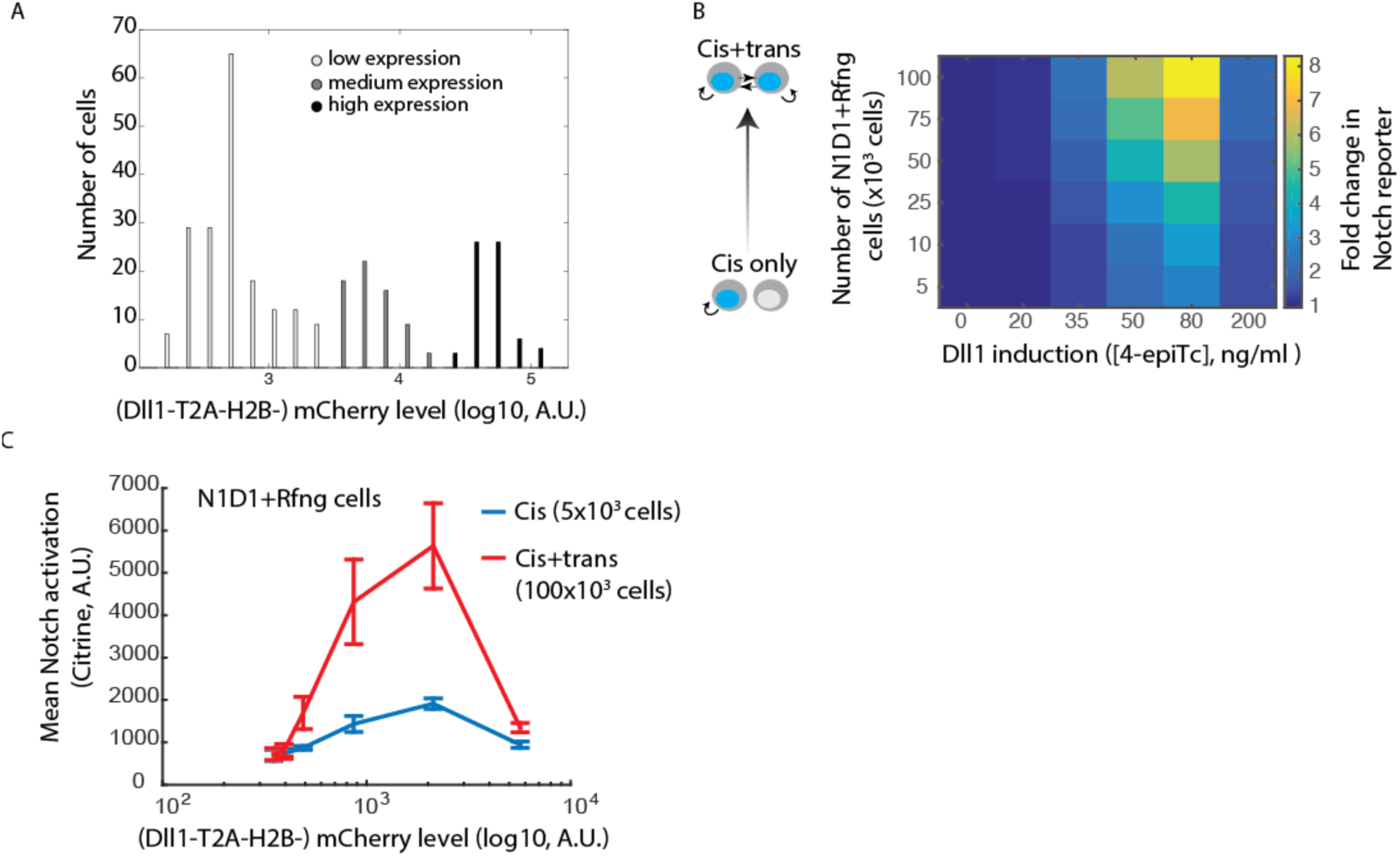
Cis- and *trans*-activation share similar features. (A) Histograms of mCherry fluorescence in cells analyzed in Figure 1E. Cells were categorized as expressing low, medium, or high Dll1 levels (shades of grey). **(B)** (*Left*) Schematic showing that total activation levels represent different relative contributions from *cis*- and *trans*-activation as the fraction of N1D1+Rfng cells increases. At the lowest fraction analyzed (lowest row of matrix, 5×10^3^ N1D1+Rfng and 150×10^3^ wild-type CHO-K1 cells), activation represents *cis*-activation, while increasing the fraction of N1D1+Rfng cells (higher rows) leads to a larger contribution of *trans*-activation to the total signal. (*Right*) Heatmap of the fold change in mean Citrine levels in N1D1+Rfng cells, relative to background Citrine levels, for a range of relative cell fractions and Dll1-induction levels. Rows defined above. Columns correspond to different concentrations of the 4-epiTc inducer. Cells were plated under the indicated conditions and analyzed less than 24 hours later by flow cytometry. Data represents 3 biological replicate experiments. **(C)** Comparison of mean activation in N1D1+Rfng cells plated at lowest (*cis*, blue line) or highest (*cis*+*trans*, red line) relative density as a function of mCherry levels, which provide a co-translational readout of Dll1 expression. Error bars represent s.e.m. of 3 biological replicates.

**Figure 1-figure supplement 3.**
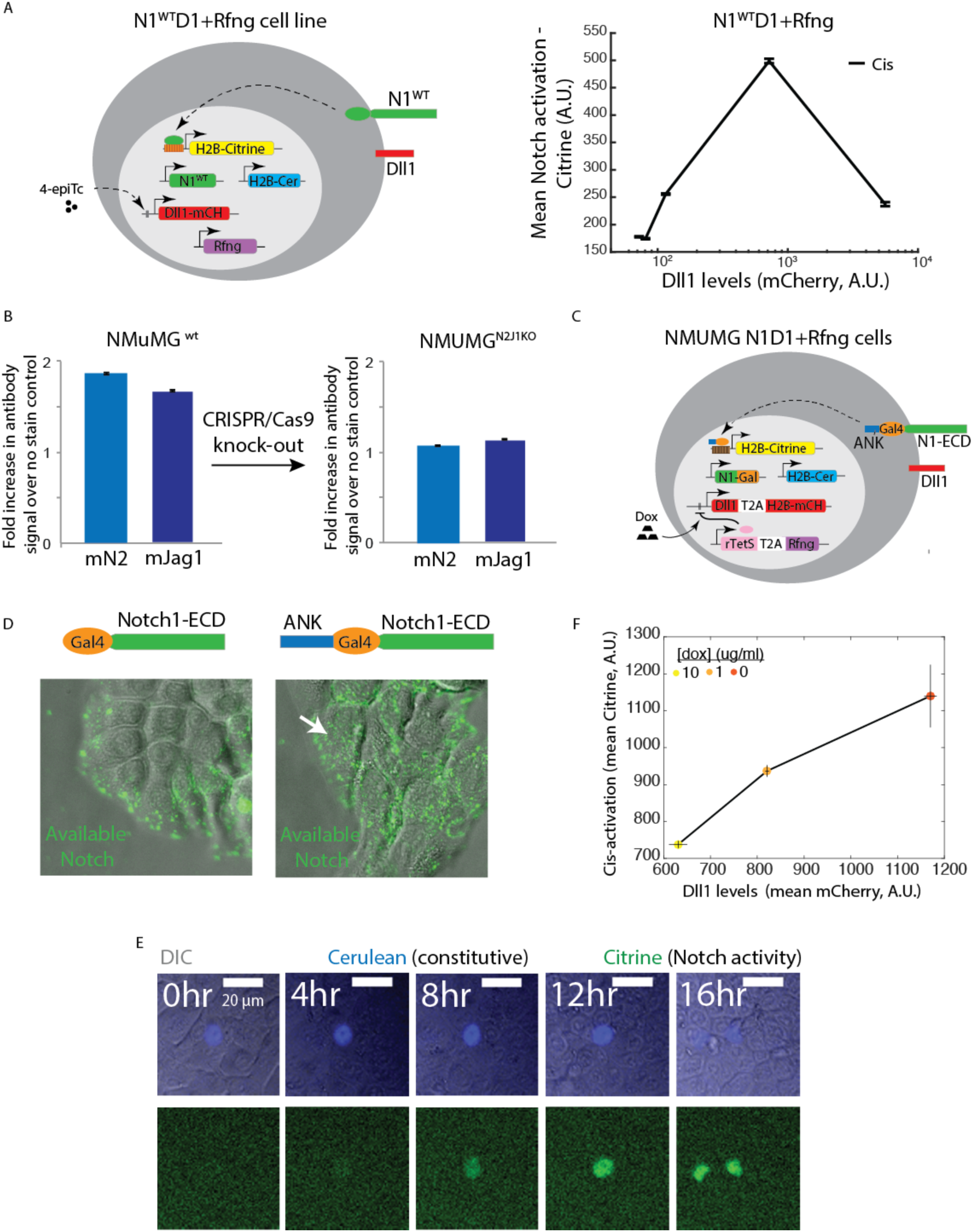
Cis-activation occurs with the wild-type Notch1 receptor and in multiple cell types. **(A)** (*Left*) The N1^WT^D1+Rfng cell line (schematic). CHO-K1 cells were engineered to express the wild-type Notch1 receptor (‘N1^WT^’, green), an H2B-Citrine reporter (yellow) activated by cleaved NICD through 12 multimerized CSL binding sites in the promoter region (orange), and a Dll1-mCherry protein (red), from a 4-epiTc inducible promoter. Constructs for constitutive expression of Rfng (purple) and H2B-Cerulean (‘H2B-Cer’, blue) were also stably integrated into these cells. (*Right*) Flow cytometry analysis of the mean activation of N1^WT^D1+Rfng cells in the *cis*-activation assay. Note non-monotonic dependence of activation on ligand levels, qualitatively similar to that observed for N1D1+Rfng cells (Figure 1), which contain the chimeric N1ECD-Gal4 receptor. **(B)** (*Left*) Antibody staining levels, relative to unstained controls, for endogenous Notch2 receptor and Jagged1 ligand in the mouse mammary epithelial cell line NMuMG. (*Right*) Staining for endogenous Notch2 and Jagged1 after CRISPR-Cas9 mediated knock-out (Materials and methods). Levels of the receptor and ligand decrease to background levels. **(C)** Schematic of the NMuMG N1D1+Rfng cell line. NMUMG^N2J1KO^ cells (panel B, right side) were engineered to express a chimeric receptor combining the Notch1 extracellular domain (‘N1ECD’, green) with the Gal4 transcription factor (orange) in place of the endogenous intracellular domain, and fused to the Ankyrin domain of the Notch1ICD (ANK, blue). When activated, Gal4-ANK is released and enables activation of a stably integrated fluorescent H2B-Citrine reporter gene (yellow) through UAS sites (brown) on the promoter. Cells also contain a stably integrated construct expressing Dll1 (red) with a co-translational (T2A, white) H2B-mCherry readout (red), from a Tet-off promoter. A constitutively expressed rTetR-HDAC4 (‘rTetS’) gene (pink) suppresses expression of the Dll1-T2A-H2B-mCherry cassette in the presence of doxycyline (‘Dox’). Rfng (purple) is expressed co-translationally with rTetS. Cells also constitutively express H2B-Cerulean (‘H2B-Cer’, blue). **(D)** Representative images showing surface staining (green) of N1ECD-Gal4 (*left*) or N1ECD-Gal4-ANK (*right*) receptors in NMuMG cells (gray overlay shows DIC channel). In the absence of the ANK domain, Notch receptor accumulates baso-laterally, and cannot be seen on the apical surface. Proper apical localization requires the ANK domain (white arrow). **(E)** Filmstrip showing activation (green depicts nuclear fluorescence in the Citrine channel) of an isolated NMuMG N1D1+Rfng cell using time-lapse microscopy. Constitutive cerulean fluorescence (blue) in the nucleus of the same cell is also shown. See Video 2 for additional examples of *cis*-activation under these conditions. **(F)** Dll1 expression levels (measured using the co-translational mCherry fluorescent protein) vs. mean activation (Citrine reporter fluorescence) of NMuMG N1D1+Rfng cells in the *cis*-activation assay. Dll1 expression was controlled by treating cells with 0, 1, or 10 ug/ml doxycycline (yellow-orange circles as indicated) (Materials and methods). Data represent the mean values across 3 biological replicate experiments, and error bars represent s.e.m.

**Figure 1-figure supplement 4.**
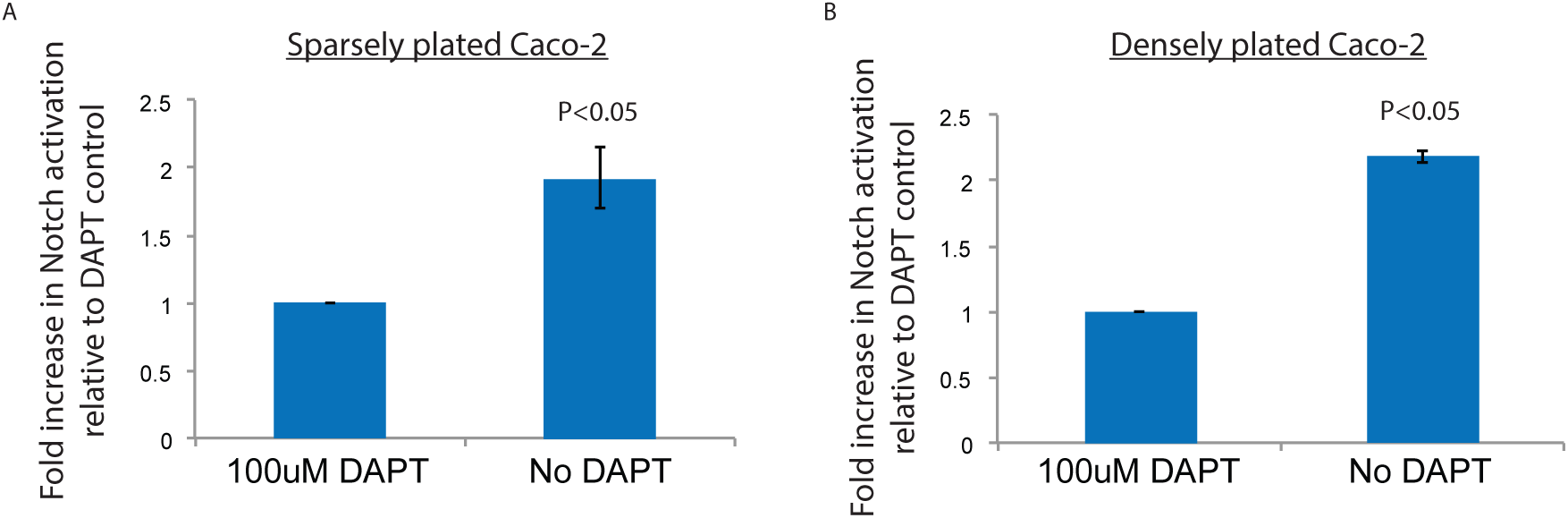
Cis-activation occurs with endogenous ligands and receptors in Caco-2 cells. **(A)** Mean activation levels in sparsely-plated Caco-2 cells, transfected with the 12xCSL-H2B-Citrine reporter construct, with or without 100 uM DAPT, <24 hours after plating, endogenous Notch activation was analyzed by flow cytometry. **(B)** Mean activation levels in densely plated Caco-2 cells, transfected with the 12xCSL-H2B-Citrine reporter construct, and treated with or without 100 uM DAPT. Endogenous Notch activation was analyzed by flow cytometry <24 hours after plating. Error bars represent s.e.m of 3 biological replicate experiments. *P*-values calculated using the one-sided Student T-test.

**Figure 2-figure supplement 1.**
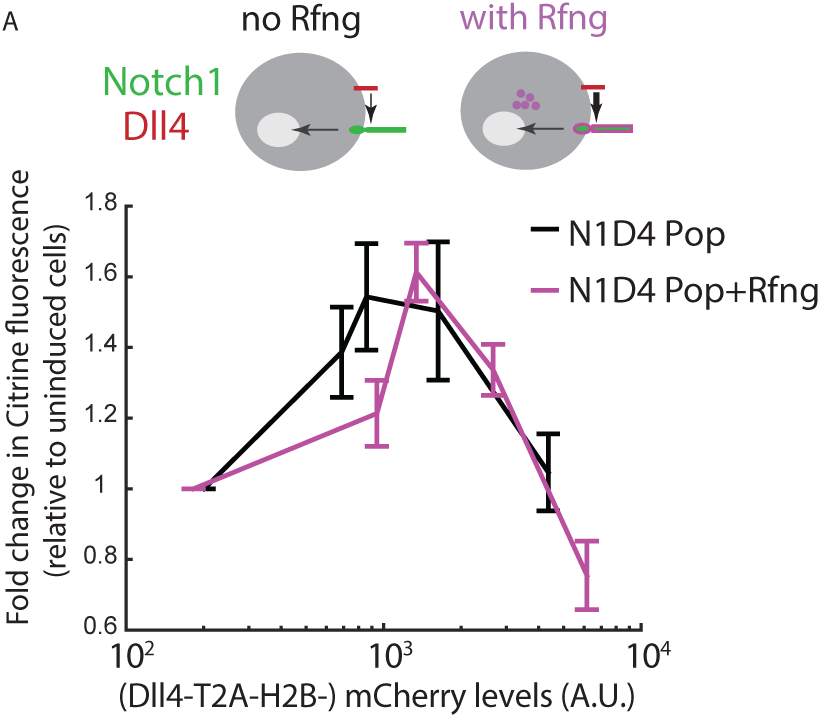
Rfng does not modify the *cis*-activation behavior of N1D4 cells. **(A)** (*Top*) Cell lines used for analyzing effect of Rfng (purple) on *cis*-activation, in the context of Notch1 Dll4. (*Bottom*) Comparison of mean *cis*-activation in polyclonal N1D4 cells with (purple, ‘N1D4 Pop+Rfng’) or without (black, ‘N1D4 Pop’) expression of Rfng, as a function of ligand expression (measured us fluorescence of the co-translated H2B-mCherry protein). Values represent mean of 3 biological replicate experiments, and error bars indicate s.e.m.

**Figure 2-figure supplement 2.**
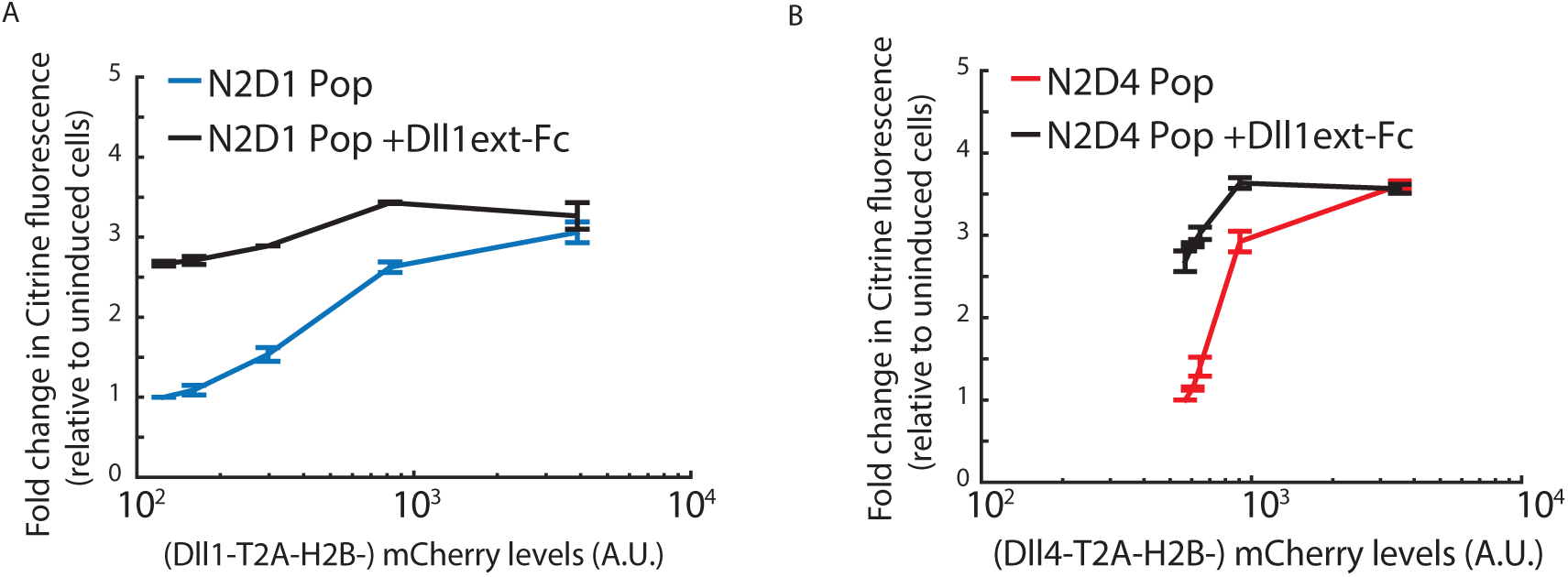
Notch2 lacks *cis*-inhibition with Dll1 or Dll4. **(A)** Mean Notch activation levels, relative to background reporter fluorescence, in polyclonal N2D1 Pop cells plated on surfaces coated with (black) or without (blue) 2.5 ug/ml recombinant human Dll1^ext^-IgG fusion protein (Materials and methods). **(B)** Mean Notch activation levels, relative to background reporter fluorescence, in polyclonal N2D4 Pop cells plated on surfaces coated with (black) or without (red) 2.5 ug/ml recombinant human Dll1^ext^-IgG fusion protein. Values represent the mean of 3 biological replicate experiments and error bars represent s.e.m. In A and B, cells were induced to express a range of Dll1/4 levels (measured using co-translated mCherry fluorescence) and cultured under *cis*-activation assay conditions (5×10^3^ N2D1/4 + 150×10^3^ CHO-K1 cells). Note similar activation levels on Dll1^ext^-IgG-coated surfaces for all *cis* Dll1/4 expression levels, suggesting that the Notch2 receptor is not inhibited by co-expressed ligand. Also note that the strength of *cis*-activation is similar to *trans*-activation by an excess of plate-bound ligand, suggesting that *cis* ligands can maximally activate Notch2-expressing cells.

**Figure 3-figure supplement 1.**
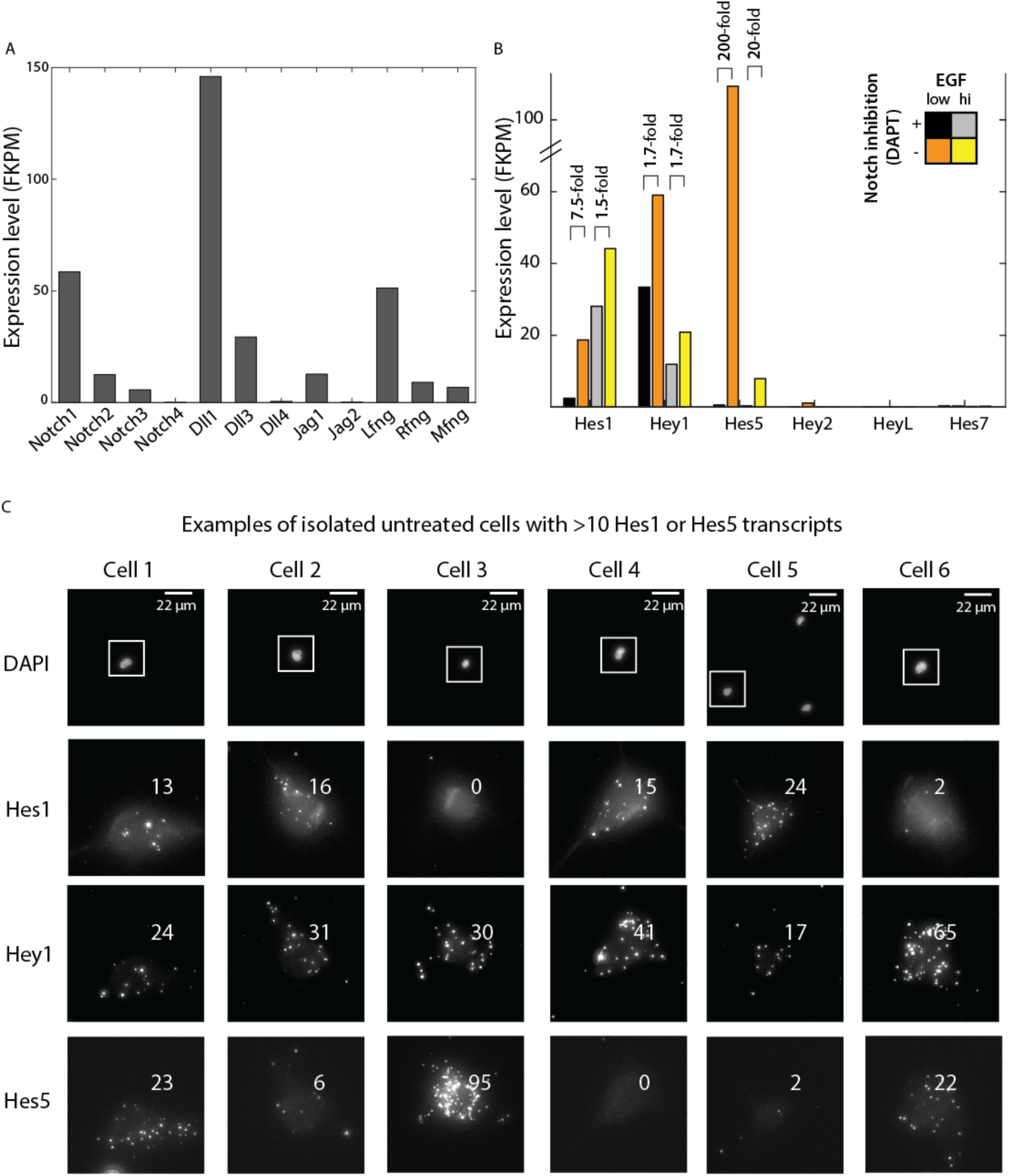
RNAseq analysis of Notch pathway component expression in neural stem cells. **(A)** Expression levels of Notch receptors, ligands, and Fringes, measured using RNAseq (see Materials and methods), in neural stem cells cultured in the presence of 0.5 ng/ml EGF and 10 μM DAPT for 12 hours. **(B)** Expression levels of canonical Notch target genes Hes1, Hey1, and Hes5 cultured in low or high growth factor conditions, with or without 10 μM DAPT treatment for 12 hours (see Materials and methods). **(C)** Examples of isolated NSCs, plated for 6 hours without DAPT, showing >10 transcripts of either Hes1 or Hes5. <3% of DAPT-treated cells pass this criterion.. Across the three target genes, the DAPT-treated sample has zero cells (out of 298 total cells) similar to Cells 1, 3, or 6, and only one cell similar to Cells 2, 4, and 5. The top row of images shows DAPI stained nuclei. Note lack of neighboring cells. Lower rows show zoomed in views of regions outlined in white, showing the cell in three fluorescent channels in which Hes1, Hey1, and Hes5 are labeled in the multiplexed smHCR-FISH experiment. Numbers indicate computationally detected number of transcripts (see Materials and methods).

**Figure 3-figure supplement 2.**
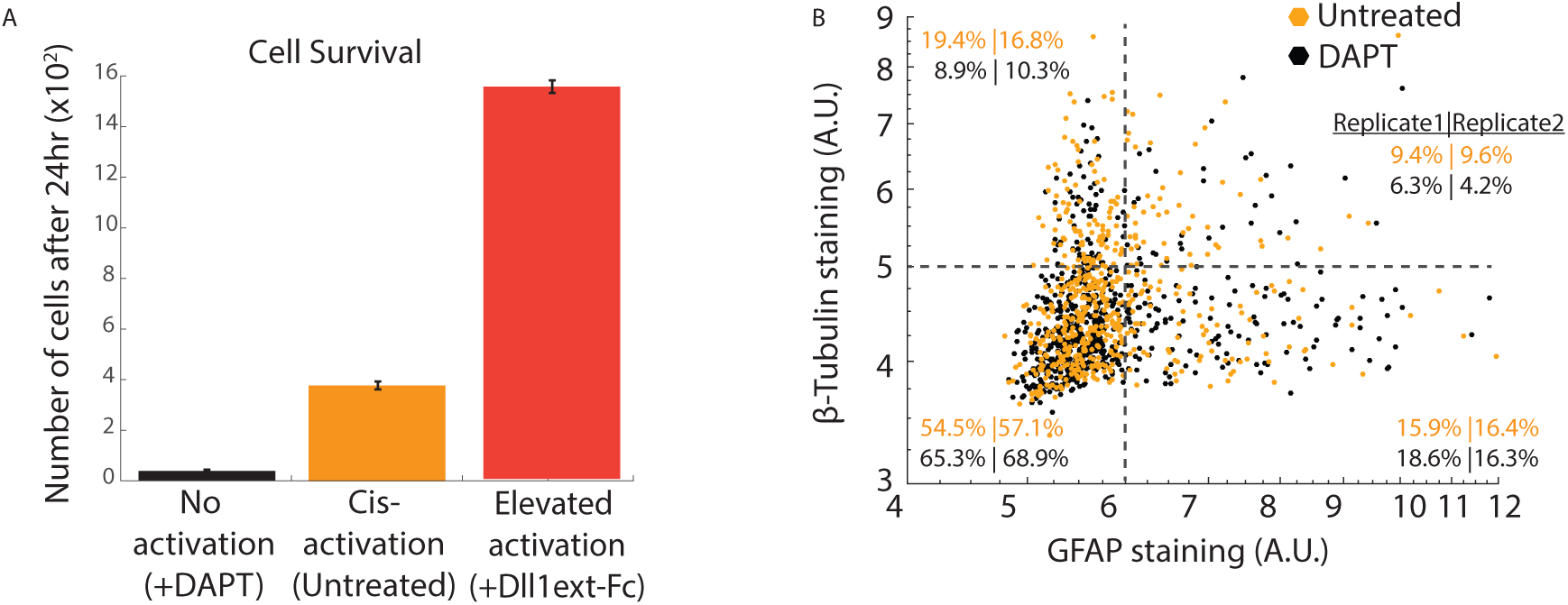
Notch-dependent cell survival and differentiation of neural stem cells. **(A)** Comparison of the number of cells after 24 hours in samples of isolated cells with (black) or without (orange) DAPT treatment, or cultured on surfaces coated with recombinant Dll1ext-IgG protein (see Materials and methods). Error bars represent s.e.m, n=4 biological replicates. **(B)** Scatter plot of immunostained GFAP and β-Tubulin in isolated cells left untreated (orange) or treated with DAPT (black), under differentiation conditions for 24 hours (see Materials and methods). Dashed lines indicate thresholds used to categorize expression as high or low, used in Figure 3D.

**Figure 4-figure supplement 1.**
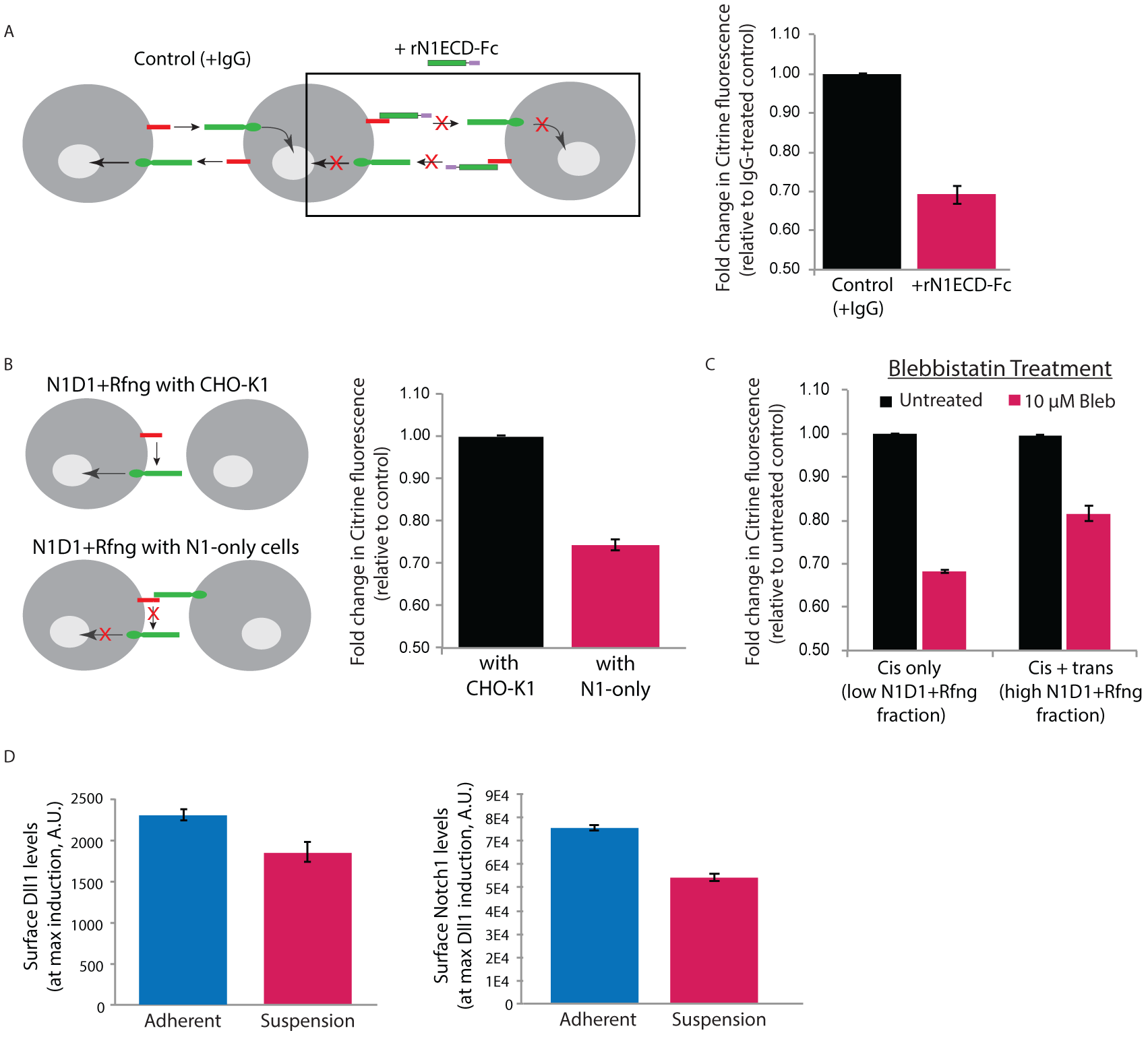
Surface perturbations affect N1D1+Rfng *cis*-activation. (A) (*Left*) Schematic showing how recombinant N1ECD-Fc protein (rN1ECD-Fc) affects *trans*-activation between cells expressing ligands (red) and receptors (green). rN1ECD-Fc protein, present in excess, binds to surface ligands and prevents their interactions with receptors on neighboring cells. This is expected to reduce overall Notch activation in cells relative to the control (cells incubated with IgG protein). (*Right*) Comparison of mean Notch activation in N1D1+Rfng cells incubated with rNotch1ECD-Fc receptors (magenta) and those that were incubated with IgG (black) for 24 hours (see Materials and methods). Cells were plated densely to allow *trans*-activation and analyzed by flow cytometry <24 hours post-plating. Error bars represent s.e.m (n=3 biological replicate experiments). **(B)** (*Left*) Schematic of N1D1+Rfng cells co-cultured with CHO-K1 cells (top) or Notch1 cells (‘N1-only cells’) that express receptor but no ligand. Notch1 cells can bind to the Dll1 ligand on the N1D1+Rfng cells and block it from interacting with Notch1 on the same cell. If surface interactions are necessary for *cis*-activation, this will lead to a decrease in Notch activation. (*Right*) Comparison of mean Notch activation in N1D1+Rfng cells co-cultured with an excess (95% of culture) of CHO-K1 cells (black) or N1-only cells (magenta). Error bars represent s.e.m (n=3 biological replicate experiments). **(C)** Mean Notch activation in N1D1+Rfng cells co-cultured at low (mostly *cis*-activation, ‘Cis-only’) or high (*cis+trans*-activation, ‘Cis + trans’) relative density with CHO-K1 cells and treated with 10 μM Blebbistatin (Materials and methods). Data is shown as a fold-change relative to control untreated samples plated similarly. Error bars represent s.e.m (n=3 biological replicate experiments). **(D)** Compared to adherent culture (blue), suspension culture (magenta) of N1D1+Rfng cells results in a slight decrease in cell-surface levels of mean Dll1 (*left*) and Notch1 (*right*), measured by immunostaining (see Materials and methods). Error bars represent s.e.m (n=3 biological replicate experiments).

**Figure 5-figure supplement 1.**
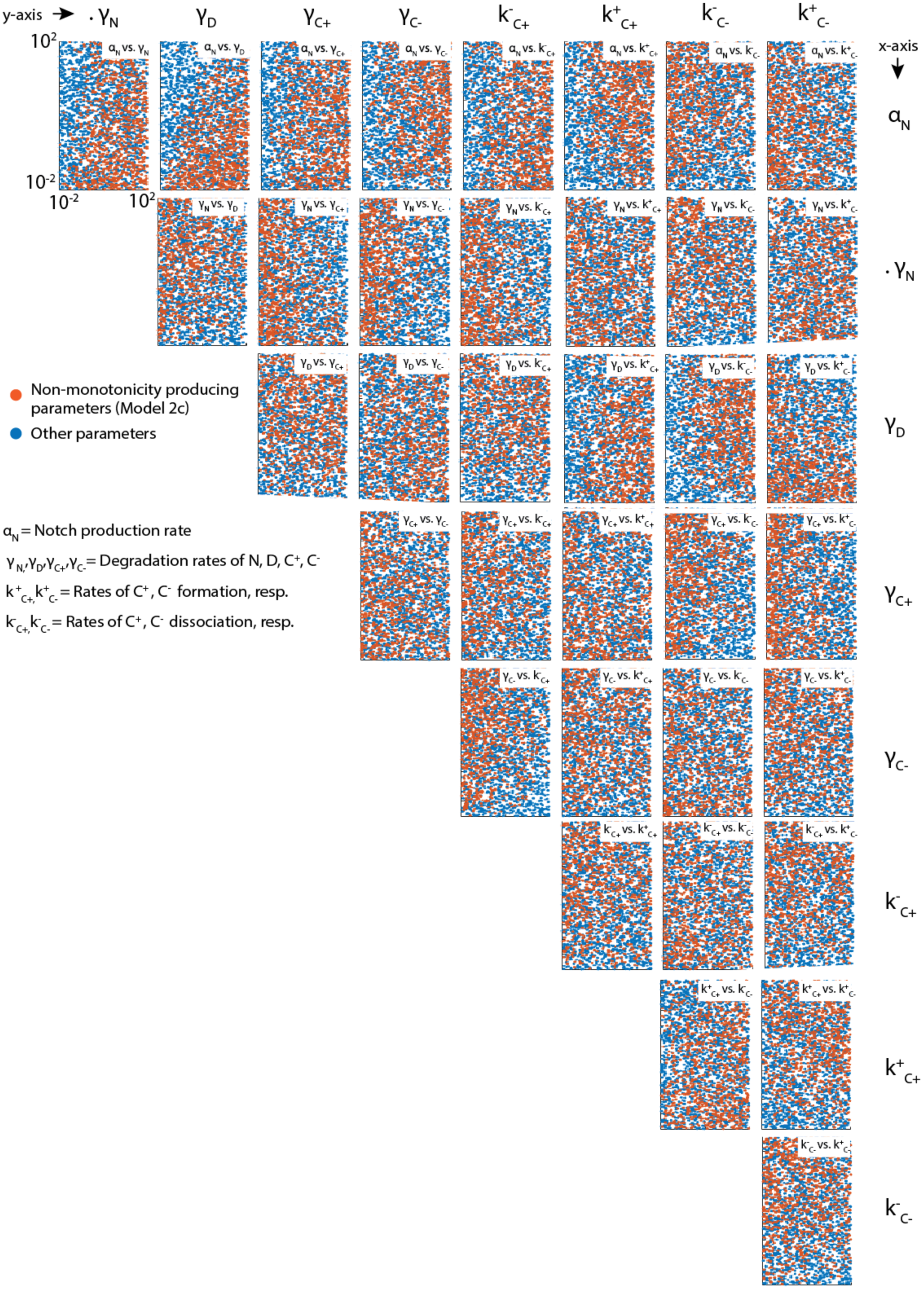
Latin Hypercube Sampling generates evenly distributed parameters. Scatter plots showing pairwise distributions of parameters tested in Models 1 and 2a-d. For clarity, each plot only shows 3000 parameter pairs, randomly subsampled from the 10,000 total parameters sets analyzed. Orange dots are subsampled from parameters that produce non-monotonic C^+^ profiles in Model 2c, while the blue dots are subsampled from the complete set of analyzed parameters. Note the even distribution of parameters across sampled space in each plot.

